# Risk assessment predicts most of the salmonellosis risk in raw chicken parts is concentrated in those few products with high-levels of high-virulent serotypes of *Salmonella*

**DOI:** 10.1101/2024.03.08.584166

**Authors:** Minho Kim, Cecil Barnett-Neefs, Ruben A. Chavez, Erin Kealey, Martin Wiedmann, Matthew J. Stasiewicz

## Abstract

*Salmonella* prevalence has reduced in U.S. raw poultry products since adopting prevalence-based *Salmonella* performance standards, but human illnesses did not reduce proportionally. We used Quantitative Microbial Risk Assessment to evaluate public health risks of raw chicken parts contaminated with different levels of all *Salmonella* and specific high- and low-virulent serotypes. Lognormal *Salmonella* level distributions were fitted using data from 2012 USDA-FSIS Baseline Survey and 2023 USDA-FSIS HACCP verification sampling data. Three different dose-response (DR) models were used: Single DR for all serotypes, reduced virulence for Kentucky, multiple serotype-specific DR models. All scenarios found risk concentrated in the few products with high *Salmonella* levels. Using a single DR model with Baseline data (μ=-3.19, σ=1.29), 68% and 37% of illnesses were attributed to the 0.7% and 0.06% of products > 1 and 10 CFU/g *Salmonella*, respectively. More recent HACCP data (μ=-4.85, σ=2.44) showed that 99.9% and 99.6% of illnesses were attributed to the 2.3% and 0.8% of products > 1 and 10 CFU/g *Salmonella*, respectively. Scenarios with serotype-specific DR models showed more concentrated risk at higher levels. Baseline data showed 91.5% and 63.7% and HACCP data showed >99.9% and 99.9% of illnesses were attributed to products > 1 and 10 CFU/g *Salmonella*, respectively. Regarding serotypes, 0.003% and 0.3% of illnesses were attributed to the 0.2% and 0.7% of products with > 1 CFU/g of Kentucky, respectively, while 69% and 78.7% of illnesses were attributed to the 0.3% and 1.2% of products > 1 CFU/g containing either Enteritidis, Infantis, or Typhimurium using Baseline or HACCP input data, respectively. These results suggest public health risk in chicken parts is concentrated in the few finished products with high-levels and specifically high-levels of high-virulent serotypes. Low-virulent serotypes, such as Kentucky, are predicted to contribute to extremely few human cases.

---

Raw poultry products are an important source of human salmonellosis cases not only in the U.S. but globally (Barrow et al., 2012). In the U.S., the United States Department of Agriculture Food Safety and Inspection Service (USDA-FSIS) developed and implemented the current performance standard based on *Salmonella* prevalence to reduce the salmonellosis cases attributable to poultry products (USDA-FSIS, 2016). USDA-FSIS data show *Salmonella* prevalence has decreased over the last several years, however this reduction in *Salmonella* prevalence has not led to the reduction of human salmonellosis cases (USDA-FSIS, 2023d; Williams et al., 2022). As outbreak-based estimations (The Interagency Food Safety Analytics Collaboration, 2023) suggest that 18.6% of foodborne salmonellosis illnesses are attributable to chicken, preventing salmonellosis from chicken products is expected to help meet the U.S. Healthy People 2030 objective of reducing *Salmonella* infections from 15.3 cases to 11.5 cases per 100,000 population (Office of Disease Prevention and Health Promotion, 2023).

Regulation based on *Salmonella* prevalence does not appear to have adequately reduced human illness cases attributed to poultry, so there is a shift toward considering the risk of high levels of contamination and different risk presented by contamination by individual serotypes, to inform potential risk management strategies (NACMCF, 2024). Studies of the *Salmonella* dose-response (DR) relationship indicated that outbreaks are often associated with higher doses causing a higher attack rate (Teunis et al., 2010). Quantitative Microbial Risk Assessment (QMRA) studies investigating *Salmonella* level-based risk management strategies suggest removing products with high-levels of *Salmonella* may substantially reduce the public health risk from chicken parts (Lambertini et al., 2019), ground turkey (Lambertini et al., 2021) and ground beef (Strickland et al., 2023).

*Salmonella* is represented by over 2,600 serotypes that can differ in their capacity to cause illness (Miller & Wiedmann, 2016). Several studies have suggested different serotypes have different likelihoods of causing illnesses when consumed in similar doses (Cheng et al., 2019; Ferrari et al., 2019; Luvsansharav et al., 2019). Outbreak data also supports that some serotypes are more commonly associated with human illnesses (Jackson et al., 2013; Jones et al., 2008). The top three most common serotypes identified in the CDC FoodNet surveillance system are Enteritidis (16% of total), Typhimurium (14%) and Newport (10%), collectively responsible for 40% of all reported salmonellosis cases from 1996-2022. Further, genomic analyses show some serotypes share common determinants of increased virulence (Fenske et al., 2023). In 2023, USDA defined three serotypes (Enteritidis, Infantis, and Typhimurium) as Key Performance Indicator (KPI) serotypes for raw poultry products for the 2022-2026 fiscal years (USDA-FSIS, 2023a). This data suggests that these three serotypes are high-virulent serotypes. This was based on the incidence of these serotypes in CDC data, their link to outbreaks, and their frequency in poultry products. In contrast, to these high-virulent serotypes, there is evidence that Kentucky may represent a low-virulence serotype. *Salmonella* Kentucky was the most frequently recovered serotype from the carcass surveillance program from USDA-FSIS, but they are less likely to cause human illnesses in the U.S. than other serotypes (Cosby et al., 2015; Richards, Kue, et al., 2023).

Concurrently, USDA-FSIS also proposed a new framework with novel strategies to control *Salmonella* in poultry products (USDA-FSIS, 2022). One of three major components is to develop new enforceable final product standards for raw poultry products focusing on *Salmonella* at certain levels and/or serotypes. In a parallel effort, USDA-FSIS proposed declaring *Salmonella* above 1 CFU/g as an adulterant in Not Ready To Eat (NRTE) breaded, stuffed chicken products (USDA-FSIS, 2023e) because NRTE breaded stuffed chicken products have repeatedly been a source of *Salmonella* outbreaks, 11 outbreaks from 1998-2022 (CDC, 2023).

This study aimed to assess the public health risk of *Salmonella* contamination of chicken parts with different levels of all serotypes and risk from specific high-virulent and low-virulent serotypes using public data collected by USDA-FSIS to define level and serotype inputs. Poultry products are good for exploring the public health impact of food contaminated with different levels and serotypes of *Salmonella* because of the large amount of contamination data available. This work advances previous risk assessment efforts by comparing results from increasingly complex DR approaches, with the most complex approach using epidemiological and human challenge data to model different serotype-specific DR relationships for four commonly recovered serotypes in chicken parts (Teunis, 2022). This work also compares results from using *Salmonella* enumeration data from a chicken parts baseline study from 2012 that was used in the previous QMRA effort (Lambertini et al., 2019), to results derived from using recent 2023 USDA-FSIS HACCP verification data for chicken parts (USDA-FSIS, 2023b). This data was obtained with qPCR, which has a much higher upper limit of quantification making it possible to better observe and model the high-level tail of the *Salmonella* level distribution.

## Material and methods

### Modeling overview

The main steps considered in the QMRA are shown in Figure 1. *Salmonella* in raw chicken parts was modeled from finished product packaging to consumption. For the *Salmonella* contamination distribution, two datasets from USDA-FSIS were collected and used (see Table 1). The QMRA focused on the assessment of the proportion of illnesses coming from products contaminated with certain levels [>LOD (=1/30 mL, equivalent to 0.0074 CFU/g), >1 CFU/g, >10 CFU/g] of all *Salmonella* or serotypes of interest including high-virulent serotypes (Enteritidis, Infantis, and Typhimurium) and a low-virulent serotype (Kentucky). The dose in one serving was calculated by multiplying values drawn from *Salmonella* level and serving size distributions. Then, a scaling factor was applied to reduce *Salmonella* levels to account for the retail and consumer between raw product packaging and consumption, such as consumer cooking. The scaling factor was set to match the average probability of illness to a baseline probability of illness, about 2 in a million, calculated using public health data. For the risk characterization, three approaches with different complexities in DR models were used to assess the public health risk from different serotypes: (1) One DR model for all serotypes; (2) Reduced virulence for Kentucky; (3) Multiple serotype-specific DR models. Each approach was used to assess and compare the public health risk residing in chicken parts contaminated with different levels of all *Salmonella* and high(Enteritidis, Infantis, Typhimurium) or low-virulent (Kentucky) serotypes. A relative risk output was defined as an expected proportion of illnesses caused by products meeting certain contamination conditions compared to all illnesses predicted from a scenario.

**Figure 1.**
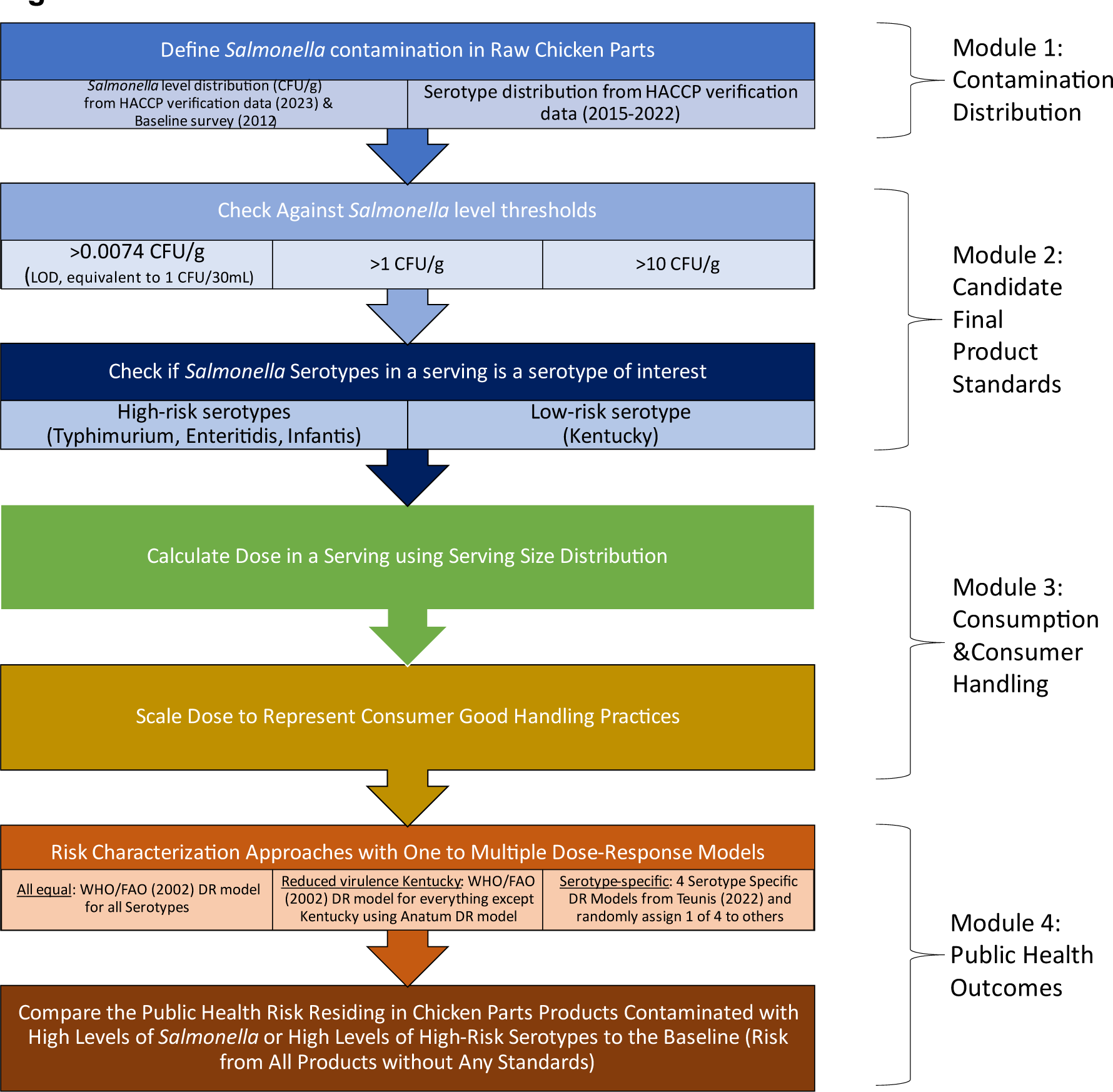
Quantitative Microbial Risk Asessment framework used in this study.

**Table 1.** Summary of collected USDA-FSIS publicly available datasets used in this project to characterize *Salmonella* contamination in chicken parts by presence, level, and serotype.

| Sampling program | Date | Available *Salmonella* relevant data | Number of records | Source |
| --- | --- | --- | --- | --- |
| The Nationwide Microbiological Baseline Data Collection Program: Raw Chicken Parts Survey (Baseline parts survey) | 01/26/2012-08/14/2012 | Screening<sup>a</sup>, serotype, level (MPN)<sup>b</sup> | 2,496 | FOIA: 2023-FSIS-00047-F<sup>d</sup> |
| USDA-FSIS routine microbiological sampling of raw chicken parts (HACCP verification data) | 01/30/2023-06/30/2023 | Screening, serotype, level (qPCR)<sup>c</sup> | 6,203 | FSIS laboratory sampling data<sup>e</sup> |
|  | 10/03/2022-01/29/2023 | Screening, serotype | 4,928 | FSIS laboratory sampling data |
|  | 03/26/2015-07/01/2022 | Screening, serotype | 70,566 | FSIS laboratory sampling data |

<sup>a</sup> *Salmonella* Screening limit of detection: 1 CFU/30 mL (0.0074 CFU/g)
<sup>b</sup> Quantifiable range for MPN enumeration method: 0.33-11 CFU/mL (0.0073 MPN/g-2.4 MPN/g)
<sup>c</sup> Quantifiable range for qPCR enumeration method: 10<sup>1</sup>-10<sup>7</sup> CFU/mL (4.5x10<sup>1</sup> CFU/g-4.5x10<sup>7</sup>CFU/g)
<sup>d</sup> https://www.fsis.usda.gov/sites/default/files/media_file/documents/FOIA-Logs_Q1-2FY23.pdf
<sup>e</sup> (USDA-FSIS, 2023b)

### Salmonella contamination data collection for chicken parts

The project obtained two different datasets including The Nationwide Microbiological Baseline Data Collection Program: Raw Chicken Parts Survey (hereafter “Baseline parts survey”) (USDA-FSIS, 2012) and USDA-FSIS routine microbiological sampling of raw chicken parts (hereafter “HACCP verification data”). Baseline parts survey data was obtained through a Freedom of Information Act (FOIA) request. HACCP verification data was obtained from the USDA-FSIS Laboratory Sampling Data webpage (USDA-FSIS, 2023b). A summary of collected data is shown in Table 1.

Three types of information were extracted from the datasets: (1) the *Salmonella* presence/absence screening test result; (2) the enumeration results based on Most Probable Number (MPN) assay in Baseline parts survey and qPCR assay (GENE-UP™ QUANT *Salmonella*) in HACCP verification data; and (3) *Salmonella* serotypes using Pulsed-Field Gel Electrophoresis (PFGE) in Baseline parts survey and Whole Genome Sequencing (WGS) in HACCP verification data. We assumed all sample collection and assays were conducted as intended, following protocols in Microbial Laboratory Guidebook (MLG) 4.14. Isolation and Identification of *Salmonella* (USDA-FSIS, 2023c). For chicken parts sampling, FSIS inspectors prepared rinsate samples by rinsing 1,814 g (4 lb) of chicken using 400 mL of Buffered Peptone Water (BPW) and sent 30 mL of rinsate sample to the lab. Then, the rinsate sample was enriched and screened through 3M™ Molecular Detection System. The primary screening result was recorded as positive or negative (LOD = 1 CFU/30 mL, equivalent to 0.0074 CFU/g) (Lambertini et al., 2019). Only samples scored positive on primary screening were used to determine the *Salmonella* level and serotypes.

### *Salmonella* contamination data fitting for level and serotype distributions

This study used a censored data analysis approach based on the limit of detection of the presence/absence screening assay (LOD = 1 CFU/30 mL) and the quantifiable range of enumeration methods (MPN: 0.33-11 MPN/mL rinse which translates to 0.007 to 2.4 CFU/g and qPCR: 10-10^7^ CFU/mL rinse which translates to 2.2-2.2x10^6^ CFU/g using the volume and weight used for the assay, respectively) to estimate the *Salmonella* level for chicken parts (USDA-FSIS, 2014). MPN assay result data were fitted to a Lognormal distribution using a Bayesian latent variable hierarchical model introduced by Williams and Ebel (2012) and as applied by Lambertini et al. (2021). This method was used to deal with uncertainty in the MPN estimates. To fit the censored data from qPCR enumeration to a Lognormal distribution, the “fitdistrplus” package was used (Delignette-Muller & Dutang, 2015). Records with negative screening results were left-censored to lower than the LOD (1 CFU/30 mL, equivalent to 0.0074 CFU/g). Records with positive screening results and <10 CFU/mL were Interval censored between LOD (1 CFU/30 mL, equivalent to 0.0074 CFU/g) and 10 CFU/mL. Observations within the qPCR quantification range (10-10,000,000 CFU/mL) were not censored. No record was right censored, as no result above 10,000,000 CFU/mL was observed.

A summary of all fits is provided in Table 2, with the observed counts from each dataset and expected proportion from fitted distribution within *Salmonella* levels of interest [> LOD (1 CFU/30 mL, equivalent to 0.0074 CFU/g), >1 CFU/g, >10 CFU/g]. The *Salmonella* level distribution fit model code is provided in GitHub, and was coded in R (R Core Team, 2023), Just Another Gibbs Sampler (JAGS) (Plummer, 2022), using the R package “rjags” package (Plummer et al., 2023). Additionally, the packages “readxl” (Wickham & Bryan, 2023), “stringr” (Wickham, 2022), “pastecs” (Grosjean & Ibanez, 2018), “knitr” (Xie, 2023), and “MASS” (Ripley & Venables, 2023) were also used in overall data management.

**Table 2.** Observed and fitted proportions of *Salmonella* in level intervals of interest.

| Counts from | Total | Negative,<br>< LOD<sup>a</sup> of<br>0.0074 CFU/g | Positive,<br>> LOD of<br>0.0074 CFU/g | Positive,<br>>1 CFU/g | Positive,<br>>10 CFU/g |
| --- | --- | --- | --- | --- | --- |
| *Baseline parts survey, data from 01/25/2012-08/13/2012* |  |  |  |  |  |
| Observed | 2,496 | 1,774<br>(71.1%) | 722<br>(28.9%) | 38<br>(1.5%) | NA<sup>b</sup> |
| Fitted Lognormal ($\mu = -3.19$, $\sigma = 1.29$) | | 79.4% | 20.6% | 0.653% | 0.0558% |
| *HACCP verification data for raw chicken parts, data from 01/30/2023 to 06/30/2023* |  |  |  |  |  |
| Observed | 6,200 | 5,661<br>(91.3%) | 539<br>(8.7%) | 56<sup>c</sup><br>(0.9%) | 25<br>(0.4%) |
| Fitted Lognormal ($\mu = -4.85$, $\sigma = 2.44$) | | 86.8% | 13.2% | 2.3% | 0.8% |

<sup>a</sup> The limit of detection (LOD) was calculated by multiplying the limit of detection of the screening assay by the volume of rinsate buffer divided by the mass of chicken: (1 MPN/30 mL rinsate) x (400 mL rinsate /1814 g parts) = 0.0074 CFU/g parts.
<sup>b</sup> Not available because of MPN method's maximum quantifiable range is 11 MPN/mL rinsate (about 2.4 CFU/g parts).
<sup>c</sup> qPCR method's quantifiable range is from 10 to 10,000,000 CFU/mL. Results reported as <10 CFU/mL values (461 counts total) were counted in the positive bin, but not for positive and >1 CFU/g or >10 CFU/g because the exact value was not known. Distribution fitting treats these censored data appropriately by interval censoring this between LOD and 10 CFU/g.

Serotype distributions from the Baseline parts survey from 2012 and two different time periods HACCP verification data (03/26/2015- 07/01/2022 and 10/03/2022-06/30/2023) are shown in Table 3. For the QMRA serotype distribution input, HACCP verification data from 2015 to 2022 was used to have more data points with a similar number of samples each year over a long recent period of time. Baseline parts survey serotype distribution was not used to avoid over-sampling from one specific year. The recently released serotype data from 10/03/2022-06/30/2023 was not used in this study because this data became available after we started using the 2015-2022 serotype distribution for the QMRA. One serotype for every serving was randomly selected from this serotype distribution to represent the serotype in one serving.

**Table 3.** *Salmonella* serotype distributions in three different datasets. A longer intermediate time HACCP verification data from 2015-2022 was used for QMRA.

| Why Included in QMRA Model | Serotype | Baseline parts survey<br>(01/26/2012-08/14/2012) |  | HACCP Verification Data<br>(03/26/2015-07/01/2022) |  | HACCP Verification Data<br>(10/03/2022-06/30/2023) |  |
| --- | --- | --- | --- | --- | --- | --- | --- |
|  |  | Count | Frequency | Count | Frequency | Count | Frequency |
| Commonly found in chicken but rare in human illnesses (Lower virulence) | Kentucky | 198 | 30.1% | 2,203 | 29.2% | 216 | 22.6% |
| Commonly associated with human illnesses in the U.S., KPI serotype<sup>b</sup> | Enteritidis | 162 | 24.6% | 1,812 | 24.0% | 229 | 24.0% |
| Commonly associated with human illnesses in the U.S., KPI serotype | Infantis | 11 | 1.7% | 1,414 | 18.7% | 301 | 31.5% |
| Commonly associated with human illnesses in the U.S., KPI serotype | Typhimurium | 60 | 9.1% | 643 | 8.5% | 108 | 11.3% |
| Association with human illness, Good dose-response data available in Teunis (2022) | Heidelberg | 50 | 7.6% | 225 | 3.0% | 2 | 0.2% |
| Non-motile Typhimurium | I 4,[5],12:i:- | 93 | 14.1% | 63 | 0.8% | 6 | 0.6% |
| All other serotypes | Other serotypes<sup>c</sup> | 84 | 12.8% | 1,195 | 15.8% | 93 | 9.7% |
| Total serotyped counts | Every serotype | 658 | 100% | 7,555 | 100% | 955 | 100% |

<sup>a</sup> For full data, see the 'Serotype data summary for parts dataset.xlsx' file in the GitHub repository.
<sup>b</sup> KPI serotype is a Key Performance Indicator serotype defined by USDA-FSIS (USDA-FSIS, 2023a)
<sup>c</sup> Other serotypes include all other serotypes as well as any samples where serotypes were unknown such as data recorded as "Multiple serotypes".

In summary, we used the Baseline parts survey and HACCP verification data to represent the different *Salmonella* level distributions, which we modeled separately to check for the impact of uncertainty in the level distribution. Conversely, we used a single serotype distribution from HACCP verification data spanning the time between both datasets used for levels as this provided a larger and more robust set of data for the serotype distribution.

### Candidate Final Product Standards

The model was designed to first check if the level is above a certain level threshold. Then it checks if the serotype in the serving is a serotype of interest or not. *Salmonella* levels of interest were set at LOD, 1 CFU/g and 10 CFU/g. LOD was set as a level of interest based on current prevalence-based *Salmonella* performance standards. 1 CFU/g was selected as a representative level for being the threshold for NRTE breaded chicken products. 10 CFU/g was set to test a higher level than 1 CFU/g.

High-virulent and low-virulent serotypes were selected based on the scientific evidence showing the relationship between human illnesses and serotypes. Table 3 lists serotypes that were considered for serotype-specific DR models in this study and the reason for that. High-virulent serotypes considered in this study were Enteritidis, Infantis, and Typhimurium because they are often related to human illnesses and USDA-FSIS also considered these three serotypes as KPIs for raw chicken products. As our risk assessment was specific for the U.S., Kentucky was considered a low-virulent serotype as it rarely causes human illness in the U.S. (few reported cases) but is one of the serotypes most commonly recovered from chicken products in the U.S. Thus, a scenario of excluding Kentucky was evaluated.

Candidate final product standards investigated for the public health risk were (1) “level only” assuming *Salmonella* above a certain level is targeted (2) “level&high-virulent (KPI) serotypes” focusing on high-levels of high-virulent *Salmonella*, (3) “level&exclude low-virulent serotype Kentucky” focusing on the high-levels of all *Salmonella* but Kentucky to see how letting Kentucky pass through the system affects the overall risk.

### Simulating *Salmonella* dose in one serving using a scaling factor to adjust for retail to consumer handling practices

The total dose of *Salmonella* per serving was calculated by multiplying observations from the *Salmonella* level distribution and the serving size distribution. For the *Salmonella* level distribution, Lognormal distribution parameters [Baseline parts survey: (μ=-3.19, σ=1.29), HACCP verification data: (μ=-4.85, σ=2.44)] were used. A previously published serving size distribution (Lambertini et al., 2019), with a mean of 2.06 Log g and a standard deviation of 0.23 Log g, was used; these data came from the NHANES data for chicken breast consumption (CDC, 2018).

A scaling factor representing multiple Log reductions was used to adjust for the fact that the input *Salmonella* distribution is raw chicken parts, but obviously people rarely consume chicken raw. Therefore, the scaling factor provides an aggregate Log reduction from finished product packaging to consumption that represents the overall effect of retail, consumer handling, and cooking practices. The value of this scaling factor was set to match the baseline mean probability of illness model output to the order of magnitude of the incidence of salmonellosis associated with chicken consumption in the U.S. The nationwide consumption of chicken was estimated based on 30.9 kg (61.8 lb) of chicken per capita consumption (USDA-FSIS, 2023). The serving size was estimated to be 114 g (as per above, 10^2.06 log g per serving) equating to around 271 servings of chicken per person in a year. When considering the U.S. population of about 332 million people on July 1^st^ 2021 (United States Census Bureau, 2023), it can then be calculated that the total chicken servings per year in the U.S. is around 90 billion servings which was calculated by multiplying the U.S. population by the number of servings per person. The number of foodborne illness attributable to chicken was calculated by multiplying a mean estimated number of 1,027,561 domestically acquired salmonellosis cases in a year from Scallan et al. (2011) by 17.3% of foodborne disease attributed to *Salmonella* in chicken from 2020 data reported in 2022 (most current number at the time of our calculation) (The Interagency Food Safety Analytics Collaboration, 2022) leading to 177,768 salmonellosis cases attributed to chicken. Using the values calculated above, the probability of illness in one serving can be obtained by dividing the amount of foodborne illness attributed to *Salmonella* in a year (177,768 cases) by the total number of chicken servings (89,969,366,913 servings) consumed. The resulting calculation produces an estimate of about 2 salmonellosis cases per 1 million servings of chicken consumed. The scaling factor for each model was calculated to match the average probability of illness to this baseline by running the model with 10 of 100,000 iterations with Monte Carlo simulation. When the mean of 10 simulations matches the baseline probability of illness (about 2 in a million), the scaling factor was saved for use in the QMRA.

### Risk characterization

For the risk characterization, three different DR model approaches with different complexities were used to assess the public health risk from different serotypes: (1) WHO/FAO (2002) DR model parameters using for all serotypes; (2) Serotype Anatum DR model parameters representing low virulent serotype Kentucky and WHO/FAO (2002) model parameters assigned for non-Kentucky serotypes; (3) Teunis (2022) four serotype-specific DR model parameters assigned to 7 serotypes and randomly assigning 1 of 4 DR model parameters to other serotypes. Table 4 summarizes which DR model parameter is assigned to which serotype and the reasoning for assigning. The average probability of illness in a serving is characterized by simulating 1 million servings.

**Table 4.** QMRA approaches for assigning available dose-response models for various *Salmonella* serotypes of interest in this project.

| Dose-Response (DR) model used for each Serotype |  |  |  |  |  |  |  |
| --- | --- | --- | --- | --- | --- | --- | --- |
| Kentucky | Enteritidis | Heidelberg | Infantis | Typhimurium | I 4,[5],12:i:- | Anatum | Others |
| QMRA approach 1. All serotypes equal |  |  |  |  |  |  |  |
| Beta-Poisson model and parameters from WHO/FAO (2002) for all serotypes |  |  |  |  |  |  |  |
| QMRA approach 2. Reduced virulence for Kentucky |  |  |  |  |  |  |  |
| Anatum DR parameters for Beta-Poisson model (Center for Advancing Microbial Risk Assessment, 2024) using a McCulloch & Elsele (1951) challenge study data to represent reduced risk of Kentucky | Beta-Poisson model with parameters from WHO/FAO (2002) for all serotypes that are not Kentucky |  |  |  |  |  |  |
| QMRA approach 3. Serotype-specific DR parameters collected from Teunis (2022)<sup>a</sup> |  |  |  |  |  |  |  |
| Anatum DR parameters were used to represent low-virulent serotype | Enteritidis specific DR parameters | Heidelberg specific DR parameters, representing high-virulent serotype and having reliable sample size (>100) | Heidelberg specific DR parameters representing high-virulent serotype because Infantis parameters did not have reliable sample size (8) | Typhimurium specific DR parameters | Typhimurium specific DR parameters because I 4,[5], 12:i:- is a non-motile Typhimurium | Anatum DR parameters | Randomly assign 1 of 4 serotype-specific DR parameters (Typhimurium, Enteritidis, Heidelberg, Anatum) |

<sup>a</sup> Hierarchical Beta-Poisson model was used [ $P(\text{inf}) + P(\text{ill}|\text{inf})$ ] following multivariate normal hyperdistribution

The probability of illness per serving was estimated using two DR models.

The first one is the Beta-Poisson DR equation (eq 1), where *S.F.* is the scaling factor, and *α* and *β* values are parameters of the distribution for calculating the probability of illness (*Pill*). Beta-Poisson model was used for all serotypes of *Salmonella* (WHO/FAO, 2002) and *Salmonella* Anatum from QMRA wiki (https://qmrawiki.org/experiments/salmonella-anatum). For Beta-Poisson DR model, Probability of illness is characterized as

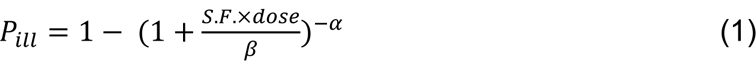

The second model is a hierarchical Beta-Poisson DR model (eq 2) for serotype-specific DR model parameters extracted from Teunis (2022). Among 14 serotype-specific DR model parameters provided, only four serotype-specific DR model parameters (Enteritidis, Heidelberg, Typhimurium and Anatum) were used here to simulate the serotype-specific probability of illness as they represented serotypes of interest and showed appropriate sample size in outbreak data (n ≥ 100 data points), and reliable human challenge studies.

Teunis (2022) serotype-specific DR models follow these two hierarchical DR functions. The probability of illness for a given dose (P*ill|dose*) is conditional (P*ill|inf*) on infection from a given dose (P*inf|dose*), as

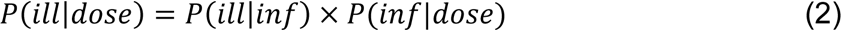

Where the probability of infection (P*inf*) is defined by the hypergeometric Beta-Poisson model (Teunis & Havelaar, 2000) using serotype-specific Teunis (2022) microbial infection parameters

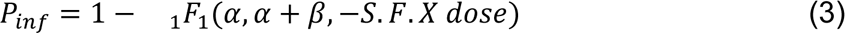

Then the probability of illness given infection uses a hazard model of illness

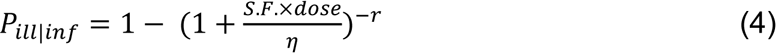

For equations 3 and 4, four parameters *α, β*, *η, r* are extracted from Teunis (2022) and provided in Table S1.

For the risk assessment, relative risk is defined as the proportion of illnesses expected from products with certain types of contamination. The proportion of finished products meeting the described contamination was also obtained from the 1 million iterations of the Monte Carlo simulation. The number of servings from products with certain levels of all *Salmonella* or serotypes of interest was calculated by multiplying the proportion by about 90 billion estimated total number of chicken servings in a year. The mean probability of illness from the products with certain levels of all *Salmonella* or serotypes of interest was then multiplied by the number of servings that are coming from the products with the contamination described. This estimated number of illnesses was then divided by the total number of illnesses expected in a year from all servings consumed. The relative risk was compared between three different QMRA approaches and two different *Salmonella* level distributions.

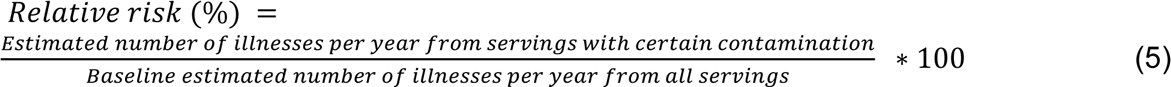

## Data and Model availability

The QMRA for this study was developed in both R and @risk. R code used for this study was developed in R version 4.1.1. (R Core Team, 2021). All QMRA models and datasets used in this study are accessible at https://github.com/foodsafetylab/Kim-2024-PoultrySalmonellaModels.

## Results

### Publicly available finished product chicken parts datasets show rare contamination with *Salmonella* above 1 CFU/g

Enumeration data from two different USDA-FSIS finished chicken parts datasets were used to compare *Salmonella* contamination at different levels (Table 2). The Baseline parts survey data from 2012 shows 722 of 2,296 (28.9%) samples tested positive using a rinse of 4 lb of parts in 400 ml of buffer, enriching and quantifying a 30 ml portion (for LOD of 1 CFU / 30 ml in rinsate, so 0.0074 CFU/g in parts). Focusing on the high-level tail, only 38 (1.52%) samples tested above 1 CFU/g, and the observed MPN results had an upper limit of 11 MPN/mL rinse, which was 2.4 CFU/g parts (so it was not possible to directly observe if samples were above 10 CFU/g). A distribution fitting process accounting for the lower detection and upper quantification limits (left and right censored data) gave a lognormal distribution with a mean of -3.19 Log(CFU/g) and a standard deviation of 1.29 Log(CFU/g). This distribution implies a 20.6% *Salmonella* prevalence (% >LOD of 0.0074 CFU/g, lower than observed), with 0.7% of samples above 1 CFU/g (close to observed), and 0.06% of the samples above 10 CFU/g.

The more recent HACCP verification data collected from Jan. to jun. 2023 shows that 539 of 6,200 (8.7%) chicken part samples tested positive. This more recent data used qPCR for quantification, not MPN as in the Baseline parts survey, which was more sensitive to the higher-level tail. For HACCP verification data, 56 (0.9%) samples tested above 2.2 CFU/g (the lower limit of quantification, lower LOQ, binned in Table 2 as > 1 CFU/g) and 25 (0.4%) tested above 10 CFU/g. These data also included 461 positive samples that tested less than the lower LOQ (binned as positive in Table 2). These samples are analyzed as interval-censored (> LOD of 0.0074 CFU/g and < lower LOQ of 2.2 CFU/g parts from the assay lower LOQ of < 10 CFU/ml rinse). A distribution process accounting for the lower detection and quantification limits (left and interval censored data) gave a Lognormal distribution with a mean of -4.85 Log(CFU/g) and a standard deviation of 2.44 Log(CFU/g). This distribution implies a 13.2% *Salmonella* prevalence (above observed), with 2.3% of samples above 1 CFU/g (close to observed), and 0.8% above 10 CFU/g (close to observed).

Across datasets, both observed counts and fitted Lognormal distribution showed that a small proportion of products had contamination above 1 CFU/g. Comparing the two datasets, the more recent HACCP verification data was estimated to have about 1 Log lower mean level distribution, but about 1 Log greater variability. These parameter estimates are consistent with the observation that the more recent HACCP verification data shows lower *Salmonella* prevalence but very likely more samples above the 1 CFU/g and 10 CFU/g high-level thresholds (differences in quantifiable range prevent direct comparison of observed frequencies at specific high-level thresholds).

### This QMRA estimates chicken parts with finished product contamination above 1 CFU/g *Salmonella* account for most of the risk, when modelling contamination data from the 2012 Baseline parts survey and a classic WHO/FAO dose response approach treating all *Salmonella* serotypes equally

Table 5 shows the proportion of total salmonellosis risk from chicken parts for those parts that match a given finished product contamination profile (the ‘relative risk’) across many different contamination profiles that could represent candidate finished product standards (rows in the table) and assumptions regarding input contamination data and DR model approaches (columns). Each result was collected with 1 million Monte Carlo iterations in R, where the mean risk of illness for the product with a given contamination profile was compared to the risk for all products (under those assumptions for contamination distribution and DR model approach), such that the baseline risk from products with all levels and all serotypes is always 100%.

**Table 5.**
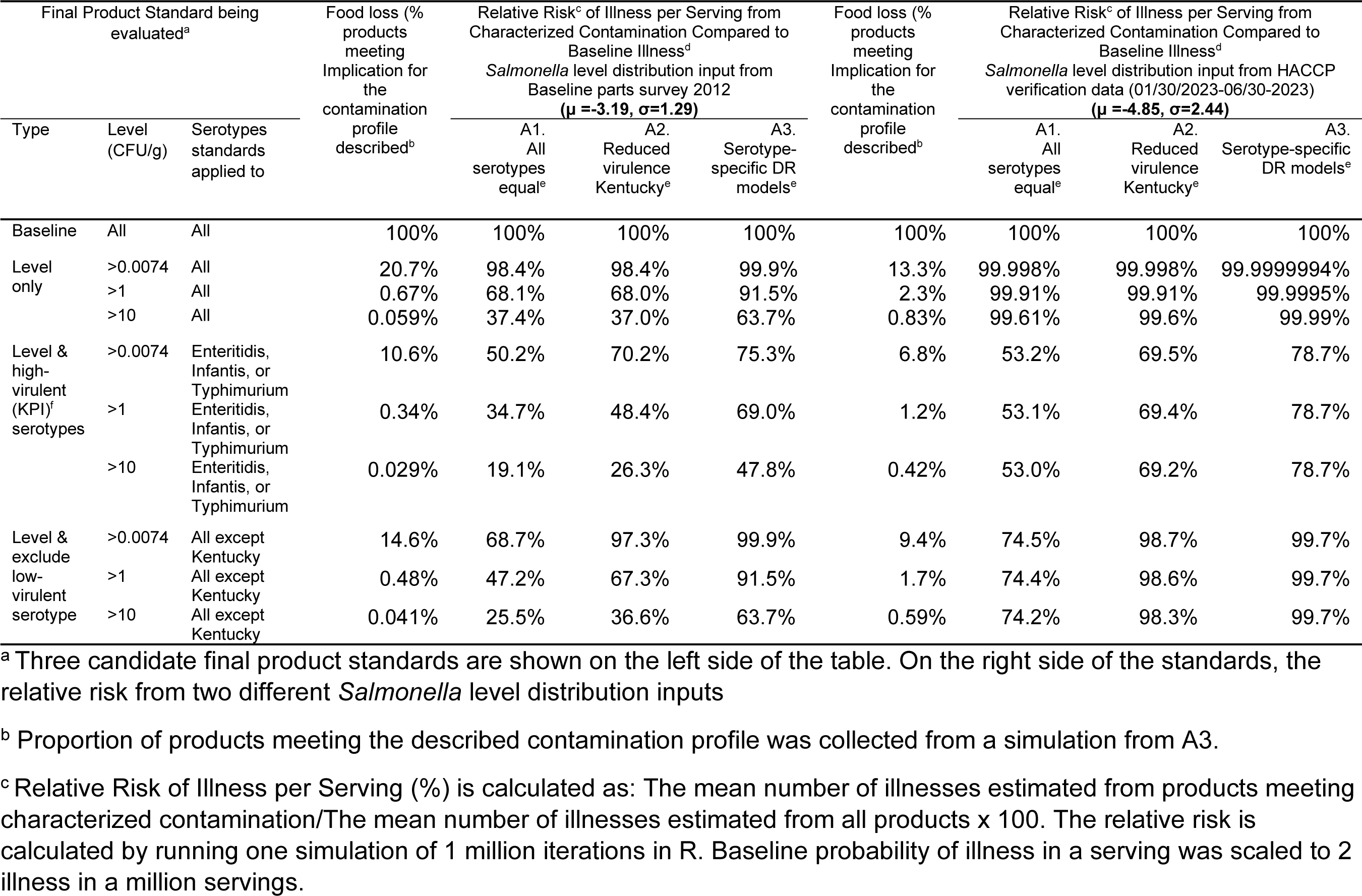
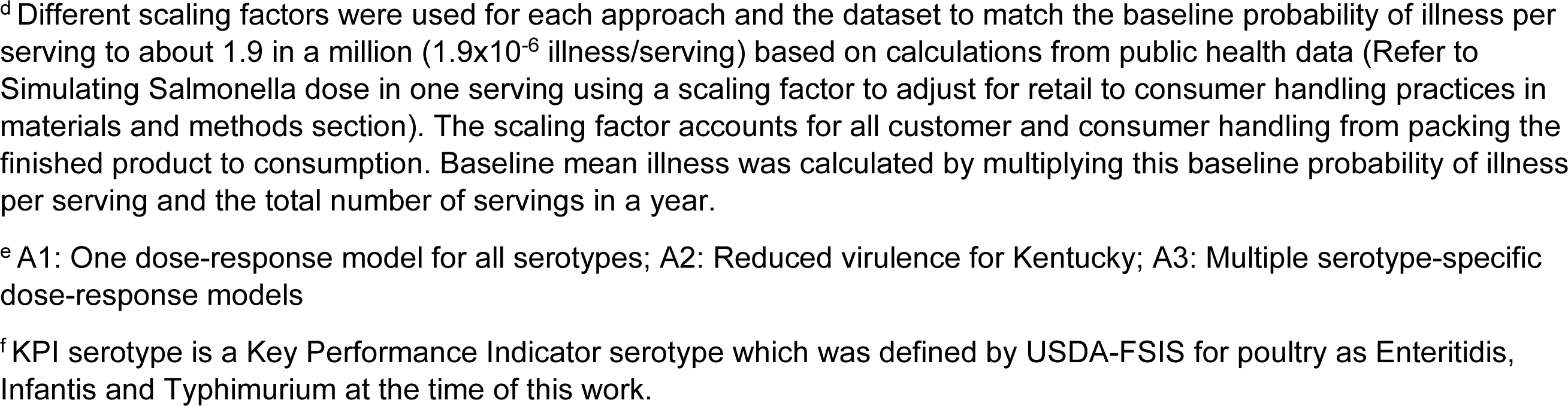
Comparing relative risk of illnesses using two different *Salmonella* distributions and three dose-response model approaches.

| Final Product Standard being evaluated<sup>a</sup> | | | Food loss (%)<br>products meeting<br>Implication for<br>the<br>contamination<br>profile<br>described<sup>b</sup> | Relative Risk<sup>c</sup> of Illness per Serving from<br>Characterized Contamination Compared to<br>Baseline Illness<sup>d</sup><br>*Salmonella* level distribution input from<br>Baseline parts survey 2012<br>($\mu = -3.19, \sigma = 1.29$) | | | Food loss (%)<br>products meeting<br>Implication for<br>the<br>contamination<br>profile<br>described<sup>b</sup> | Relative Risk<sup>c</sup> of Illness per Serving from<br>Characterized Contamination Compared to<br>Baseline Illness<sup>d</sup><br>*Salmonella* level distribution input from HACCP<br>verification data (01/30/2023-06/30-2023)<br>($\mu = -4.85, \sigma = 2.44$) | | |
| --- | --- | --- | --- | --- | --- | --- | --- | --- | --- | --- |
| Type | Level<br>(CFU/g) | Serotypes<br>standards<br>applied to |  | A1.<br>All<br>serotypes<br>equal<sup>e</sup> | A2.<br>Reduced<br>virulence<br>Kentucky<sup>e</sup> | A3.<br>Serotype-specific DR<br>models<sup>e</sup> |  | A1.<br>All<br>serotypes<br>equal<sup>e</sup> | A2.<br>Reduced<br>virulence<br>Kentucky<sup>e</sup> | A3.<br>Serotype-specific<br>DR models<sup>e</sup> |
| Baseline | All | All | 100% | 100% | 100% | 100% | 100% | 100% | 100% | 100% |
| Level<br>only | >0.0074 | All | 20.7% | 98.4% | 98.4% | 99.9% | 13.3% | 99.998% | 99.998% | 99.9999994% |
|  | >1 | All | 0.67% | 68.1% | 68.0% | 91.5% | 2.3% | 99.91% | 99.91% | 99.9995% |
|  | >10 | All | 0.059% | 37.4% | 37.0% | 63.7% | 0.83% | 99.61% | 99.6% | 99.99% |
| Level &<br>high-virulent<br>(KPI)<sup>f</sup><br>serotypes | >0.0074 | Enteritidis,<br>Infantis, or<br>Typhimurium | 10.6% | 50.2% | 70.2% | 75.3% | 6.8% | 53.2% | 69.5% | 78.7% |
|  | >1 | Enteritidis,<br>Infantis, or<br>Typhimurium | 0.34% | 34.7% | 48.4% | 69.0% | 1.2% | 53.1% | 69.4% | 78.7% |
|  | >10 | Enteritidis,<br>Infantis, or<br>Typhimurium | 0.029% | 19.1% | 26.3% | 47.8% | 0.42% | 53.0% | 69.2% | 78.7% |
| Level &<br>exclude<br>low-virulent<br>serotype | >0.0074 | All except<br>Kentucky | 14.6% | 68.7% | 97.3% | 99.9% | 9.4% | 74.5% | 98.7% | 99.7% |
|  | >1 | All except<br>Kentucky | 0.48% | 47.2% | 67.3% | 91.5% | 1.7% | 74.4% | 98.6% | 99.7% |
|  | >10 | All except<br>Kentucky | 0.041% | 25.5% | 36.6% | 63.7% | 0.59% | 74.2% | 98.3% | 99.7% |

<sup>a</sup> Three candidate final product standards are shown on the left side of the table. On the right side of the standards, the relative risk from two different *Salmonella* level distribution inputs
<sup>b</sup> Proportion of products meeting the described contamination profile was collected from a simulation from A3.
<sup>c</sup> Relative Risk of Illness per Serving (%) is calculated as: The mean number of illnesses estimated from products meeting characterized contamination/The mean number of illnesses estimated from all products x 100. The relative risk is calculated by running one simulation of 1 million iterations in R. Baseline probability of illness in a serving was scaled to 2 illness in a million servings.
<sup>d</sup> Different scaling factors were used for each approach and the dataset to match the baseline probability of illness per serving to about 1.9 in a million ( $1.9 \times 10^{-6}$ illness/serving) based on calculations from public health data (Refer to Simulating Salmonella dose in one serving using a scaling factor to adjust for retail to consumer handling practices in materials and methods section). The scaling factor accounts for all customer and consumer handling from packing the finished product to consumption. Baseline mean illness was calculated by multiplying this baseline probability of illness per serving and the total number of servings in a year.
<sup>e</sup> A1: One dose-response model for all serotypes; A2: Reduced virulence for Kentucky; A3: Multiple serotype-specific dose-response models
<sup>f</sup> KPI serotype is a Key Performance Indicator serotype which was defined by USDA-FSIS for poultry as Enteritidis, Infantis and Typhimurium at the time of this work.

The most appropriate starting point to present results is those assuming contamination from the Baseline parts survey and using the WHO/FAO(2002) DR model approach that treats all serotypes equally (4^th^ and 5^th^ columns), as these assumptions are most consistent with previous risk assessments. For these assumptions, 98.4% of illnesses were attributed to the 20.7% of finished chicken parts that have *Salmonella* levels above the typical assay LOD of 0.0074 CFU/g. Further, 68.1% of illnesses were attributed to the 0.7% of finished products above 1 CFU/g and 37.4% of illnesses were attributed to the 0.06% of finished products above 10 CFU/g. Therefore, under these assumptions, most of the illnesses from chicken parts are predicted to be from finished products contaminated with more than 1 CFU/g.

### When accounting for serotype-specific dose-response models, the estimated relative risk of salmonellosis caused by finished products above 1 CFU/g increases to almost all the total risk

The next two DR model approaches model increasingly complex treatment of serotype-specific risk, first reducing the virulence of Kentucky (Table 5, 6^th^ column), then using 4 different serotype-specific DR parameters sets to represent the range of *Salmonella* serotypes (Table 5, 7^th^ column). Simply reducing Kentucky virulence has almost no impact on the relative risk of products contaminated with high-levels of any *Salmonella.* Accounting more fully for serotype-specific virulence noticeably increases the relative risk associated with high-level contamination, from 68.1% to 91.5% for products above 1 CFU/g and from 37.4% to 63.7% for products above 10 CFU/g, still using the Baseline parts survey contamination. The QMRA also predicts a small increase in illness risk associated with the 20.7% of products with detectable contamination, from 98.4% to 99.8%.

Having a serotype-specific DR model makes it possible to examine the relative risk presented by various serotypes and levels of contamination. The bar plots at top of Figure 2 visualize these relative risks for products above the level threshold on the right side of dashed line, breaking these apart by serotype groups of interest, and also products below the level threshold on the left side. The density plots below the bar plots visualize the proportion of finished products that are above or below the level thresholds for both input contamination datasets. The results that assume the Baseline parts survey contamination, show that the single grouping with the most illness attributed (75.3%) is the three KPI serotypes (Enteritidis, Infantis, and Typhimurium) above the given level thresholds. *Salmonella* Kentucky is predicted to be responsible for relatively few illnesses, even when above the relatively high-level thresholds.

**Figure 2.**
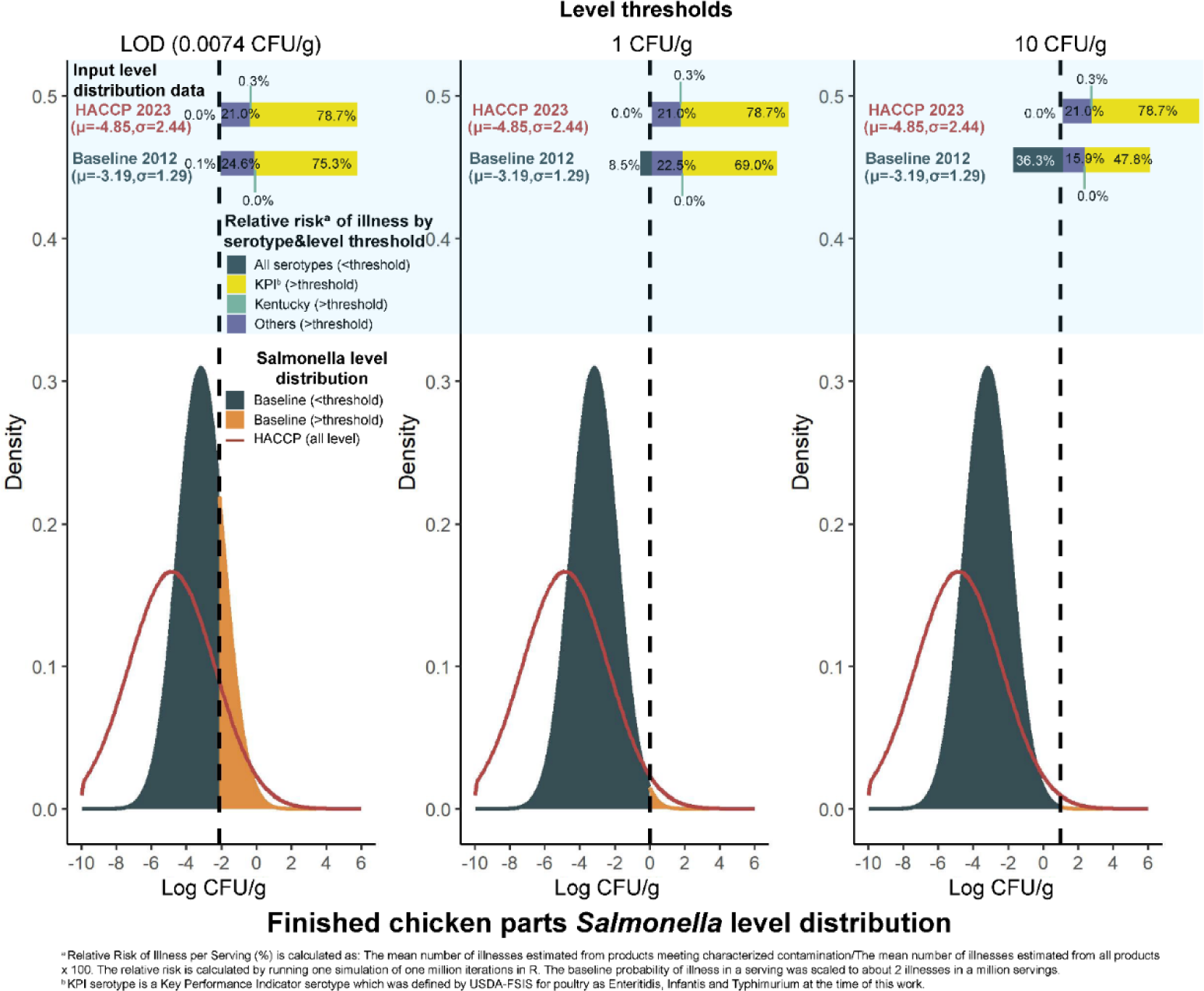
Comparison of the relative risk of illnesses attributed to various *Salmonella* serotype groups above or below given level thresholds. Two *Salmonella* level distribution inputs were used, and all results represented are from the serotype-specific dose-response approach (A3). The colored layer at the above part of the figure shows the relative risk coming from each *Salmonella* level distribution input. The density functions below visualize *Salmonella* level distributions from the two different datasets used. The dashed line separates both risk and level distribution into below and above the level threshold. The relative risk from different serotypes is divided into three high-virulent KPI serotypes (Enteritidis, Infantis, and Typhimurium), a low-virulent serotype Kentucky, and other serotypes, each shown as different colors.

### When assuming contamination consistent with more recent chicken parts HACCP verification data, the higher though still rare fraction of products with very high-level contamination (0.8% above 10 CFU/g) is estimated to contain virtually all of the public health risk

The more recent HACCP verification data shows a lower prevalence, but larger tail of high-level contamination, which is modeled as a contamination input distribution with a lower mean but higher deviation (Table 5, last 4 columns, and Figure 2). Using these data, virtually all the risk (99.61%) is attributable to the 0.8% of products above 10 CFU/g, using the WHO/FAO DR model approach treating all serotypes equally. Risk is even more concentrated (99.99% or more) in products above 10 CFU/g using the serotype-specific DR model approach. Products with detectable *Salmonella* level now represent more than 99.998% of risk across all DR model approaches. Further, 78.7% of total risk is attributable to the 0.4% of products with three KPI serotypes when present above the 10 CFU/g level thresholds (Figure 2).

### Final product standards that target relatively high-levels of high-virulent serotypes would target finished products responsible for a large fraction of public health risk while implicating a smaller fraction of finished products

Figure 2 shows that most of the public health risk is in the relatively small fraction of finished products contaminated above 1 CFU/g or 10 CFU/g, and particularly those contaminated with high-virulent serotypes including Enteritidis, Infantis, and Typhimurium across both finished product contamination datasets. Therefore, we modeled the relative risk of products that would be implicated by candidate finished products standards that apply not just to levels (e.g., > 10 CFU/g) but also what serotypes the standards would apply to (e.g., current KPI serotypes).

One analysis was a finished product standard that applies to only selected high-virulent serotypes (specifically Enteritidis, Infantis, and Typhimurium as the current USDA-FSIS KPI serotypes as of late 2023, middle rows of Table 5). Modelling the contamination from the Baseline parts survey, and using the full serotype-specific DR approach, 75.3%, 69.0%, and 47.8% of relative risk was attributed to products contaminated with detectable, >1, and >10 CFU/g respectively. The more interesting thing is the comparison between these results and the level-only standard. While 99.9% of relative risk is present in the 20.7% of products with detectable *Salmonella*, 75.3% of the total risk is present in the only 10.6% of products with detectable current KPI serotypes, suggesting most of the risk is in about half of the detectably contaminated products. Similarly, while 91.5% of the risk is in 0.6% of products with contamination >1 CFU/g of all serotypes, 69.0% of the risk is in the 0.3% of products with KPI serotypes >1 CFU/g. Furthermore, 63.7% of the risk is in the 0.06% contaminated by all serotypes >10 CFU/g and 47.8% of the risk is in the 0.03% contaminated by KPI serotypes >10 CFU/g. In all cases, standards that target only KPI serotypes would implicate about half the product and retain more than half the public health benefit, suggesting a path to effective illness reduction that meets the 25% Healthy People 2023 target.

When modelling the newer HACCP verification data, the overall trend is still present that risk can be more efficiently reduced by targeting specific serotypes, although the specifics differ. In these results, level-only standards would remove almost all the public health risk (99.99+%), but targeting only the current KPI serotypes would remove about 78.7% of relative risk while still implicating only about half the overall product. It is also worth noting that the same standards apply much less effectively when assuming the WHO/FAO(2002) DR approach treating all serotypes equally, which is a logical consequence of the DR model and further justification for work to use serotype-specific DR models for risk assessment.

As a logical converse to the above analyses targeting high-virulent serotypes, we also modeled standards that permit the presence of a low virulent serotype (specifically allowing Kentucky at any contamination level, bottom rows of Table 5). Applying this finished product standards approach to the contamination representing the Baseline parts survey, the relative risk was 99.9%, 91.5%, and 63.7%, for contamination detectable, >1, or > 10 CFU/g, respectively. Critically, these numbers are essentially the same as for the only level-based standards and do implicate about 30% less product; the same trend appears in the results for the DR approach that explicitly reduces Kentucky virulence from the single WHO/FAO DR base approach. For the contamination profile representing the newer 2023 HACCP verification data, these standards implicate products responsible for 99.7% of relative risk, compared to 99.99% or more for only level-based standards. Overall, these analyses suggest the finished products contaminated with serotype Kentucky present a very low public health risk.

## Discussion

### Because multiple risk assessments show the most risk is from high-levels of contamination, it is important to accurately measure and model the high-level tail of the contamination distribution

This and another recent chicken parts QMRA (Lambertini et al., 2019) show that the risk is concentrated in products with the high-levels of *Salmonella* (products >1 MPN/g and >10 MPN/g), as do studies in ground turkey (Lambertini et al., 2021; Sampedro et al., 2024) and ground beef (Strickland et al., 2023). But all these studies model product contamination data collected using MPN method for the enumeration (in our case the Baseline parts survey data), which unfortunately has an upper limit of quantification lower than levels responsible for much of the risk (assay limit of 11 MPN/ml, here parts rinse, which translates to 2.4 CFU/g parts). So, it is not possible to directly observe, e.g., contamination at or above 10 CFU/g.

The current best practice for building a risk assessment accounting for the unobserved higher-level tail of contamination has been to fit an underlying statistical distribution to these ‘censored’ data such that the fitted distribution gives results that would reasonably reproduce the observed pattern of both quantifiable and outside the quantification limit data. Our work uses a Bayesian latent variable hierarchical model suggested by Williams and Ebel (2012) specifically for MPN data, and was previously applied in the risk assessments by Lambertini et al. (2019, 2021). This method reasonably assumes the observed data comes from an underlying Lognormal distribution, which is commonly used for modelling chicken product contamination distributions (Jongenburger et al., 2015). The method also gives more precise and less biased parameter estimates than Maximum Likelihood Estimation (Williams & Ebel, 2012). Our analysis fitting the censored data predicts approximately 0.06% of chicken parts represented in the Baseline parts survey data would have contamination > 10 CFU/g, but one would have much greater confidence in this estimate of the tail of the distribution if one could directly observe contamination at that level.

One major advance in our work is that this is the first chicken risk assessment (to our knowledge) that uses public *Salmonella* data for chicken parts contamination that directly measures contamination in the high-level tail. Specifically, our study evaluates 2023 HACCP verification data which uses qPCR for enumeration, a method with higher and wider quantification range than MPN (qPCR: 10 to 10^7^ CFU/mL rinse compared to MPN: 0.33 to 11 MPN/mL rinse, which translates to qPCR 2.2 to 2.2x10^6^ CFU/g parts compared to MPN 0.007 to 2.4 CFU/g parts). The modest tradeoff between these methods is that when using qPCR in combination with the limit of quantification (2.2 to 2.2x10^6^ CFU/g parts) is wider than for MPN (0.007 to 2.4 CFU/g parts). An even more sensitive method, such as MPN-qPCR-Shortened Incubation Time (SIT) introduced by Kim et al. (2017), could possibly resolve the tradeoff between quantifying the highest levels and those just above those detected by enrichment. Still, censored data analysis can accommodate any range of ‘interval censored’ data. The input 2023 HACCP verification data had 0.4% of samples >10 CFU/g, and the fitted distribution had 0.8% of contamination >10 CFU/g, which is a reasonable fit to the now observable highest risk tail, while still reproducing the prevalence in the data reasonably (observed 8.7% positive, fitted 13.2% above the detection threshold). In contrast, the older 2012 Baseline parts survey data showed a higher overall prevalence (observed 28.9%, fitted 20.6%), with a small tail of high-level contamination (1.5% observed > 1 CFU/g, fitted as 0.6%, with no ability to observe > 10 CFU/g, fitted as 0.06%). It seems to be a valuable advance in data to support risk assessment to directly measure the highest levels, and to give direct support to the distribution fitting processes to model those levels, given that risk becomes increasingly concentrated at higher levels of contamination. Such effort to more accurately model high-level tails is very important in this case, where the more recent data suggests a decrease in overall prevalence, but indicates a possible increase in the high-level contamination tail.

### High-virulent serotypes are commonly found in finished product samples, so it is important to accurately assess risk using serotype-specific dose-response models

There is substantial evidence that the >2,600 *Salmonella* serotypes can differ in virulence and public health risk (Fierer & Guiney, 2001; Grimont & Weill, 2007; Jones et al., 2008) with clear genomic and epidemiologic data that isolates representing some serotypes and clades show a substantially lower or higher risk of causing human illness (Fenske et al., 2023; Teunis, 2022). However, no previously published poultry QMRA adequately modeled risk comparing factors including prevalence and level with a comprehensive treatment of serotype-specific DR model (NACMCF, 2024).

Our study used three different DR approaches to illustrate the implications and importance of serotype-specific DR models. In our simplest approach, treating all serotypes as equally virulent (with contamination consistent with the Baseline parts survey data), about half of the risk in products with high-levels (>10 CFU/g, which represents about 37% of the total risk) was from those with high-levels of high-virulent serotypes (19% of the 37% total) because three high-virulent serotypes account for about half of serotype prevalence in chicken parts as you can see in 4^th^ column of Table 3. In the slightly more complex approach where we explicitly lowered the virulence of Kentucky, about two-thirds of the risk from high-level products was from high-virulent serotypes (26% of the 37% total). Two other risk assessments used a conceptually similar approach, assigning reduced virulence DR parameters to low-virulent serotypes though not just Kentucky, and reported a similar result that most illnesses were attributed to high-virulent serotypes in turkey (Sampedro et al., 2024) and beef (Strickland et al., 2023) products.

This study is the first risk assessment for chicken combining impacts of high-level contamination with a more complete serotype-specific DR approach. Specifically, the four most commonly recovered serotypes from chickens (Kentucky, Enteritidis, Infantis and Typhimurium), as well as a few others where data were available, were assigned the DR model parameters from Teunis (2022), which used epidemiological outbreak data (for high-virulent serotypes) and human feeding trials data (for low-virulent serotypes). We also adopted the source work’s approach to separately model the probability of infection and illness if infected, and the correlation between parameters. Using this more complete serotype-specific DR approach (and assuming *Salmonella* input level from the Baseline parts survey data), indicates that about two-thirds (67%) of the total risk was in products with high-levels (>10 CFU/g), and still about two-thirds of that risk was attributable to the current KPI serotypes >10 CFU/g (48% of the 67% total), with virtually no risk coming from Kentucky. This trend is even more extreme in the scenario assuming the 2023 HACCP verification data *Salmonella* input level distribution, with a larger high-level tail, where virtually all the risk is in the high-level tails, and 79% of the total risk is attributed to KPI serotypes. In this case, still almost no risk (0.3%) is attributable to Kentucky, even though this serotype represents about 30% of the contamination at all level thresholds.

One limitation in this serotype-specific risk assessment approach is that we assumed only one serotype exists in any given product. We took this approach as the current HACCP verification data only reports a single serotype for a sample, due to methodological limitations. A new method suggested by Thompson et al. (2018) can identify multiple serotypes from one sample using sequencing of CRISPR spacers present in the sample; a recent study using this method found that only 3 out of 38 post-chill carcass samples contained multiple serotypes (Richards, Siceloff, et al., 2023). In addition, the same study found that Kentucky and the three KPI serotypes (Enteritidis, Infantis and Typhimurium) were still the most frequently identified serotypes, suggesting the assumption of a single serotype for contamination will not bias these serotype-specific model results in any obvious direction.

### These results suggest one could efficiently manage food safety risk using strategies that target the small fraction of the highest risk products

As previously discussed, our QMRA (and others) showed concentrated risk in the small fraction of products contaminated with high-levels of *Salmonella*, and even more concentrated risk in products with high-levels of high-virulent serotypes. These risk assessment results imply that it would be both effective and efficient to develop and implement practical risk management strategies focusing on reducing the small fraction of high-level *Salmonella* contamination, and/or specifically high-virulent serotypes.

There are different ways to interpret this risk assessment result. First, if one assumes perfect sampling and testing could identify and eliminate every product with contamination above a given threshold, our results assess the risk reduction expected from eliminating that contamination from the food supply. Obviously, perfect sampling and testing are neither realistic nor practical, but these results could then inform a plausible maximum effect of a test and reject strategy. Other risk assessments have extensively modeled the impacts of different within lot variability (Lambertini et al., 2019; Lambertini et al., 2021), and different sampling and testing plan assumptions (Sampedro et al., 2024), which can be seen as a way to put these ‘perfect test’ results into context.

Another way to interpret this risk assessment results is to use improved process controls (Cohn et al., 2023) to dramatically reduce the likelihood of a high-level and/or high-virulent serotype product, which could then provide a meaningful level of risk reduction. One process control strategy could be adopting control limits for total *Salmonella* levels, such as setting an internal upper control limit to something observable and actionable (e.g., >1 CFU/g), and setting an upper specification limit to something indicative of a true public health failure (e.g., >10 CFU/g). Such a process control approach would allow for routine monitoring and process improvement through corrective action, without relying on a lot-specific microbial criterion to identify failed lots. The proposed USDA-FSIS *Salmonella* framework does mention the consideration of applying Statistical Process Control (SPC) to indicator organism testing throughout the sanitary dress process (USDA-FSIS, 2022). In addition, a paper has explored SPC for various quality indicators in processing facilities in the European Union (Mataragas et al., 2012) and a few recent papers have explored statistical relationships between indicator organisms and *Salmonella* at different processing stages. (Chavez-Velado et al., 2024; Williams et al., 2015; Williams et al., 2017), but none directly apply SPC to pathogens directly.

Another process control strategy can be serotype-specific control strategies. One commonly used example is the vaccination of live birds in poultry houses against one or a few high-virulent serotypes, e.g., Typhimurium (Dórea et al., 2010; NACMCF, 2024; Hofacre et al., 2021). The U.S. has seen a substantial decrease in infections caused by Typhimurium and Heidelberg last 20 years and this seems to match with the timing when commercial poultry vaccines against Typhimurium became available (NACMCF, 2024). Another example could be preharvest lotting strategies, where one would test incoming flocks for high-virulent serotypes and apply additional risk reduction measures to those flocks. This may include removal of high-virulent flocks from certain secondary processing, such as grinding, to reduce the risk profile of finished products from those processes. Alternatively, ‘logistical slaughter’ could be used where high-virulent flocks are processed at the end of a shift to prevent cross contamination to lower risk and is already implemented in many countries in EU (Rasschaert et al., 2020). However, the effectiveness of this approach is unclear yet as the correlation between the preharvest and finished product is shown to be weak (Nauta et al., 2009; Rasschaert et al., 2008). More specifically, there is a shift in the recovered serotype distribution between breeder flocks, chilled carcasses, and finished intact parts (Siceloff et al., 2022), complicating the relationship between preharvest interventions and finished product outcomes. In addition, vaccination targeting specific serotypes can create a niche and this niche can be filled by other serotypes, possibly other high-virulent serotypes (Foley et al., 2011).

### A strategy to target risk concentrated in contamination with high-levels or high-virulent serotypes can be applied to other chicken products and other meat and poultry

This study used the chicken parts category as a representative product because it is a widely consumed product with extensive contamination data available. Our results are consistent with a previous study (Lambertini et al., 2019), which also showed that finished products with high-level contamination have more risk, a concept relevant to other poultry and other meat products. Lambertini et al. (2021) modeled ground turkey and suggested a similar result, that a microbial criterion of 1 cell/g for finished products would lead to an estimated 46.1% reduction in risk of illness. Another QMRA using ground turkey data showed removing ground turkey lots contaminated with above 10 MPN/g and 1 MPN/g will reduce illness about 38.2% and 73.1%, respectively (Sampedro et al., 2024). A QMRA for ground beef showed a small fraction of high-level products contaminated with above 10 MPN/g and 1 MPN/g contain about 13.6% and 36.7% of risks, respectively (Strickland et al., 2023). Overall, our data and previous studies across products thus support that products contaminated with a small fraction of higher level *Salmonella* have high-virulent.

Importantly, while our study used *Salmonella* as a proxy to define *Salmonella* subgroups that differ in virulence, it is becoming increasingly clear that there is substantial diversity and heterogeneity within many *Salmonella* serotypes. This is particularly apparent and important for *Salmonella* Kentucky, which represents at least two distinct clades, one that is virulence attenuated and common in chickens in the US (representing predominantly ST 152) and one that appears highly virulent and often is resistant to multiple antibiotics (ST 198) (Tate et al., 2022), and potentially other clades when using different nomenclature (Richards, Kue, et al., 2023). In our study here, *Salmonella* Kentucky thus represents a proxy for *Salmonella* Kentucky ST 152. In such instances, future risk assessments will benefit from approaches that consider evolutionary-supported definitions of *Salmonella* clades that differ in virulence [for example, as defined by WGS or Multilocus Sequence Typing (MLST)].

## Conclusions

Contamination with high-levels of high-virulent serotypes is relatively rare in finished chicken parts. Yet this risk assessment suggests that most of the public health risk from chicken parts is concentrated in those rare products with high-levels of high-virulent serotypes. This conclusion is consistent with multiple previous risk assessments (Lambertini et al., 2019; Lambertini et al., 2021; Sampedro et al., 2024) for individual finished poultry products that build more detailed models of potentially confounding factors like lot-to-lot variation and imperfect testing. Our work advances these previous efforts by incorporating a serotype-specific DR approach based on epidemiological data and using more recent HACCP verification data that directly measure the high-level tail of the *Salmonella* contamination distribution. Therefore, this study supports a growing consensus that public health could be improved by *Salmonella* risk management strategies targeted to reduce high-levels of high-virulent serotypes so that the poultry industry is appropriately incentivized to manage the *Salmonella* risk in finished products by reducing the highest risk outcomes. In addition, our study supports that limited public health benefits would be gained from the control of low virulent serotypes (such as the virulence attenuated *Salmonella* Kentucky clade circulating in the U.S.) (Tate et al., 2022). This is important as regulatory policies centered on *Salmonella* prevalence alone may implicitly or explicitly encourage reductions of virulence attenuated *Salmonella*, such as Kentucky. This may not only not show measurable public health outcomes, but may have negative consequences, such as facilitating increased prevalence and emergence of other, possibly high-virulent *Salmonella* serotypes.

## Acknowledgments

This study was supported by a US Poultry and Egg Association grant to Stasiewicz and Wiedmann, for Project #BRF-015 Risk Assessment Comparing Alternative Approaches to Regulating *Salmonella* in Poultry by Public Health Impact Factors. In addition, both Stasiewicz and Wiedmann are members of a Coalition for Poultry Safety Reform which includes volunteer members from the poultry industry, consumer groups, and academics.

## Supplemental Material

**Table S1.**
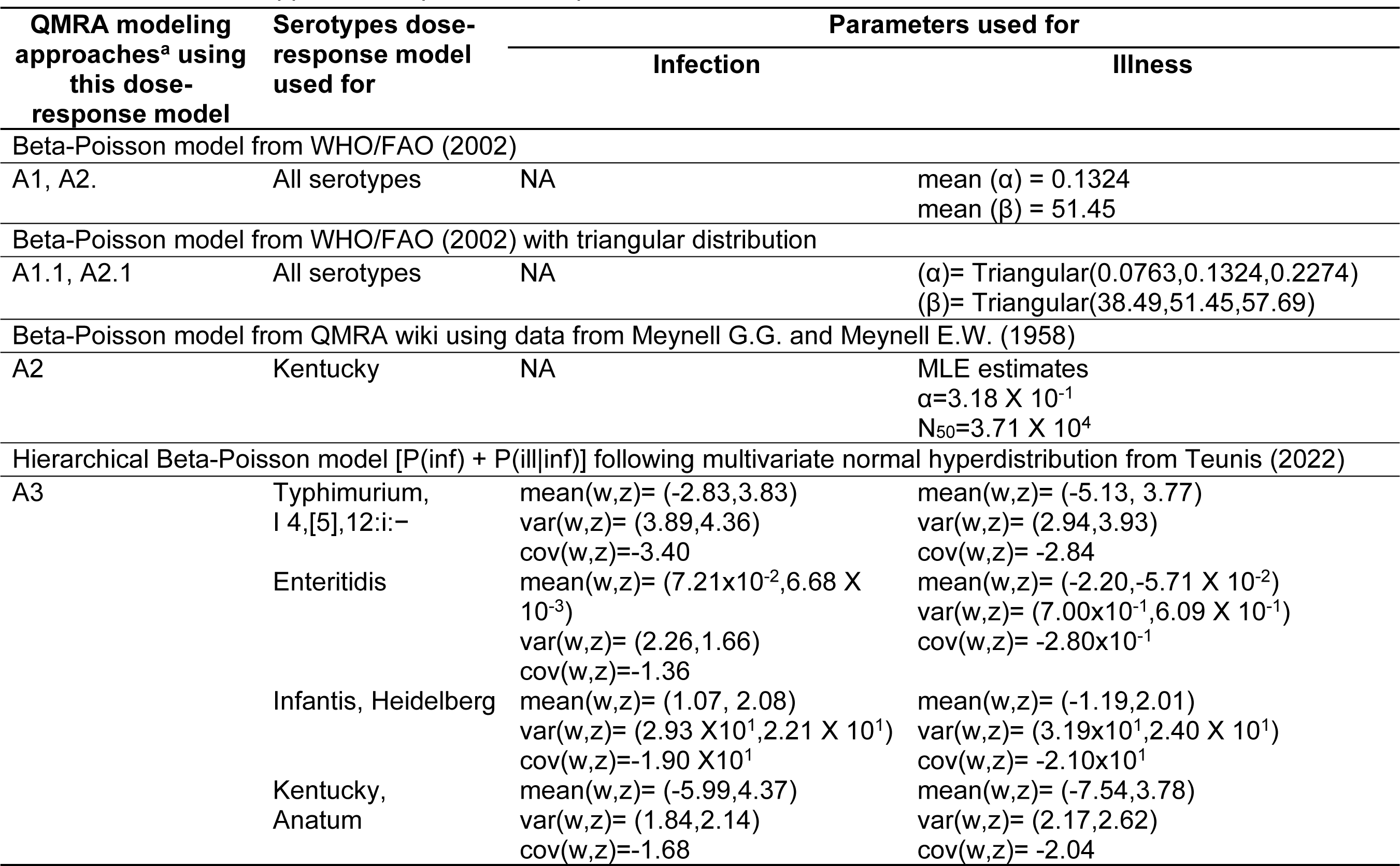
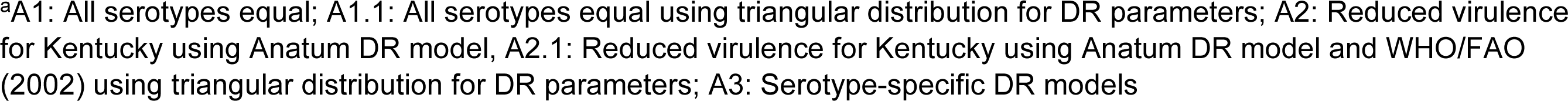
*Salmonella* spp. dose-response model parameters used in the QMRA.

| QMRA modeling approaches<sup>a</sup> using this dose-response model | Serotypes dose-response model used for | Parameters used for |  |
| --- | --- | --- | --- |
|  |  | Infection | Illness |
| Beta-Poisson model from WHO/FAO (2002) |  |  |  |
| A1, A2. | All serotypes | NA | mean ($\alpha$) = 0.1324<br>mean ($\beta$) = 51.45 |
| Beta-Poisson model from WHO/FAO (2002) with triangular distribution |  |  |  |
| A1.1, A2.1 | All serotypes | NA | ($\alpha$)= Triangular(0.0763,0.1324,0.2274)<br>($\beta$)= Triangular(38.49,51.45,57.69) |
| Beta-Poisson model from QMRA wiki using data from Meynell G.G. and Meynell E.W. (1958) |  |  |  |
| A2 | Kentucky | NA | MLE estimates<br>$\alpha$=3.18 X 10<sup>-1</sup><br>N<sub>50</sub>=3.71 X 10<sup>4</sup> |
| Hierarchical Beta-Poisson model [P(inf) + P(ill\|inf)] following multivariate normal hyperdistribution from Teunis (2022) |  |  |  |
| A3 | Typhimurium, I 4,[5],12:i:- | mean(w,z)= (-2.83,3.83)<br>var(w,z)= (3.89,4.36)<br>cov(w,z)=-3.40 | mean(w,z)= (-5.13, 3.77)<br>var(w,z)= (2.94,3.93)<br>cov(w,z)= -2.84 |
|  | Enteritidis | mean(w,z)= (7.21x10<sup>-2</sup>,6.68 X 10<sup>-3</sup>)<br>var(w,z)= (2.26,1.66)<br>cov(w,z)=-1.36 | mean(w,z)= (-2.20,-5.71 X 10<sup>-2</sup>)<br>var(w,z)= (7.00x10<sup>-1</sup>,6.09 X 10<sup>-1</sup>)<br>cov(w,z)= -2.80x10<sup>-1</sup> |
|  | Infantis, Heidelberg | mean(w,z)= (1.07, 2.08)<br>var(w,z)= (2.93 X10<sup>1</sup>,2.21 X 10<sup>1</sup>)<br>cov(w,z)=-1.90 X10<sup>1</sup> | mean(w,z)= (-1.19,2.01)<br>var(w,z)= (3.19x10<sup>1</sup>,2.40 X 10<sup>1</sup>)<br>cov(w,z)= -2.10x10<sup>1</sup> |
|  | Kentucky, Anatum | mean(w,z)= (-5.99,4.37)<br>var(w,z)= (1.84,2.14)<br>cov(w,z)=-1.68 | mean(w,z)= (-7.54,3.78)<br>var(w,z)= (2.17,2.62)<br>cov(w,z)= -2.04 |

<sup>a</sup>A1: All serotypes equal; A1.1: All serotypes equal using triangular distribution for DR parameters; A2: Reduced virulence for Kentucky using Anatum DR model, A2.1: Reduced virulence for Kentucky using Anatum DR model and WHO/FAO (2002) using triangular distribution for DR parameters; A3: Serotype-specific DR models

**Table S2.**
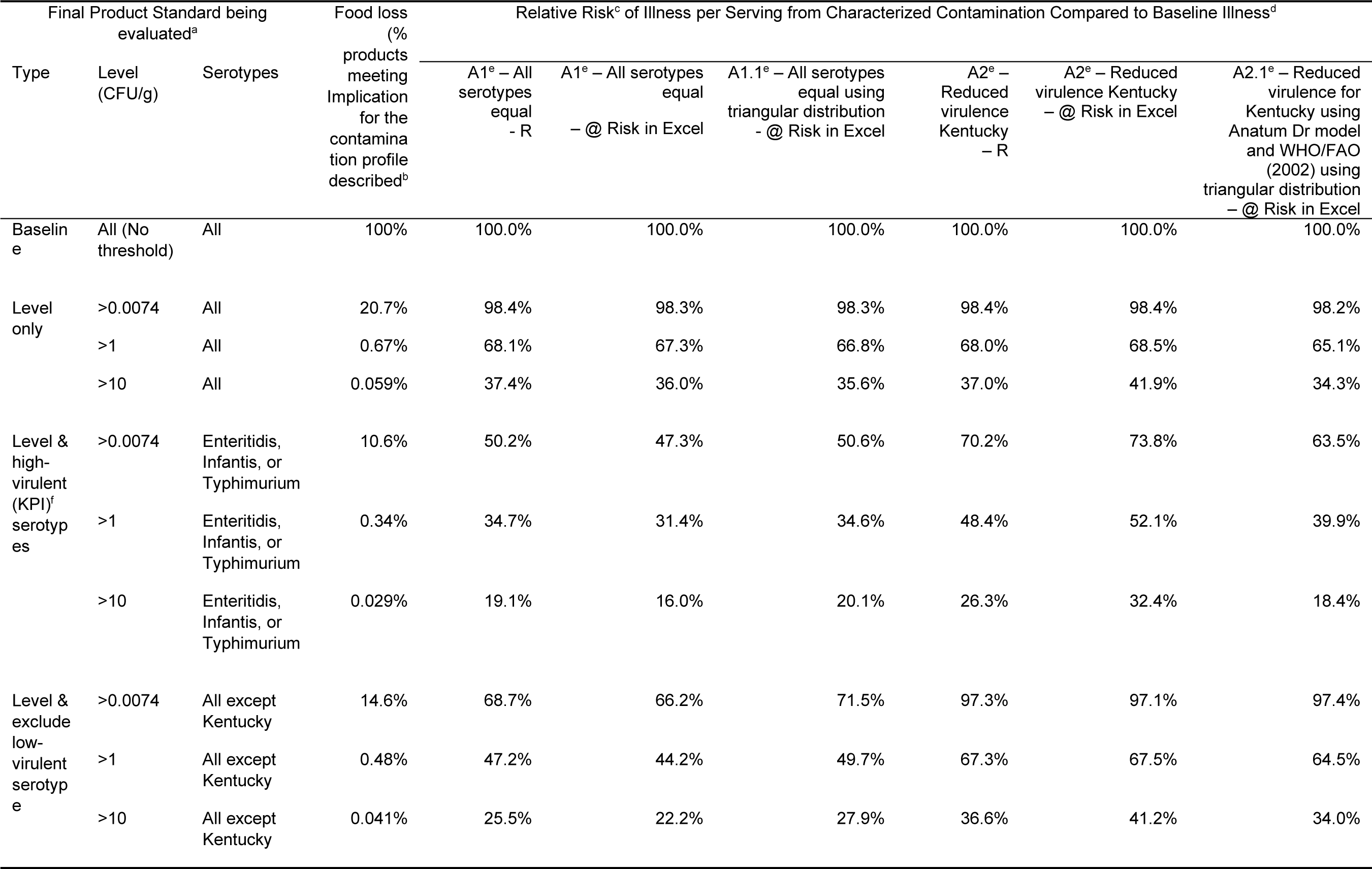
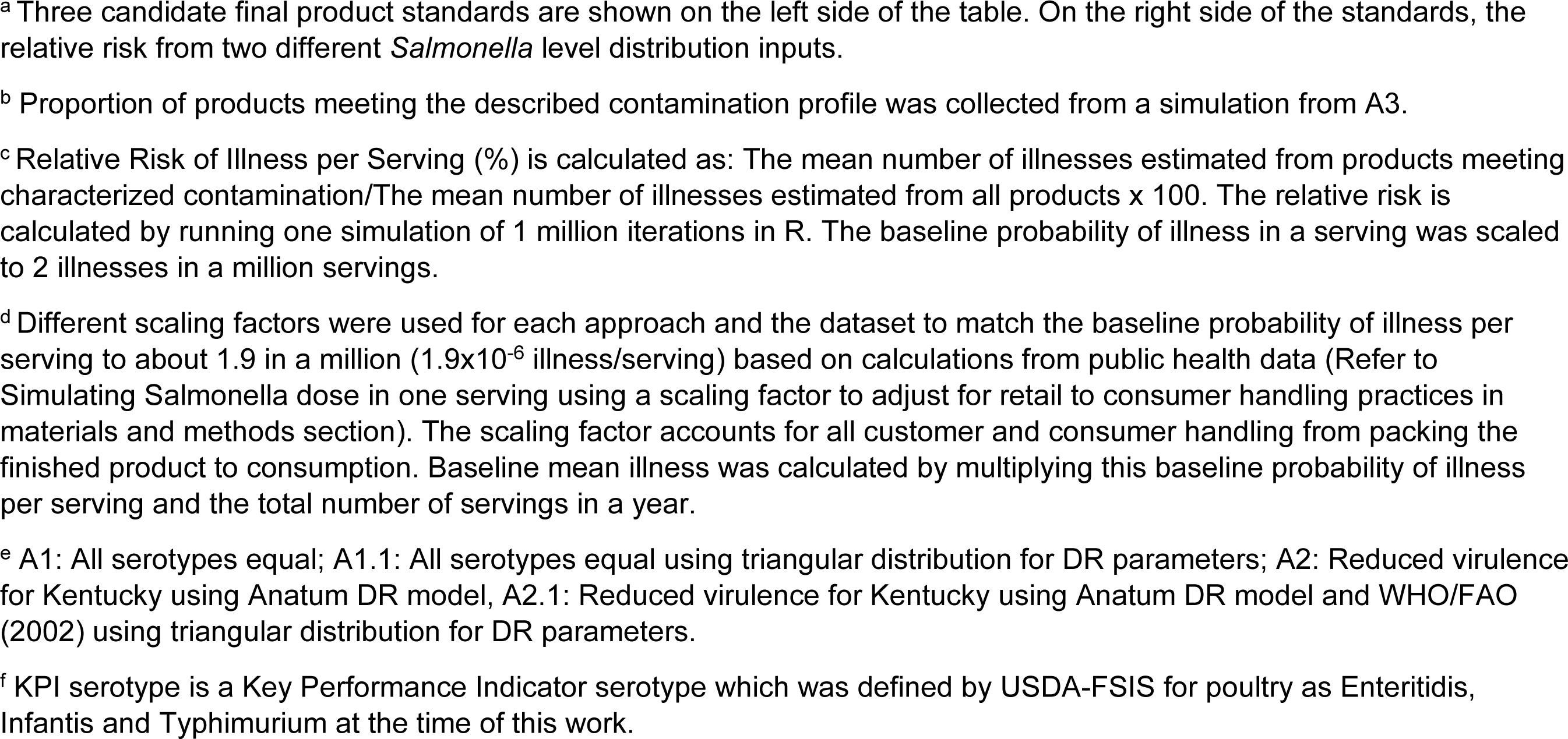
Comparing relative risk of illnesses for the implementation of models in R or @RISK for Excel from Baseline parts survey input level data.

| Final Product Standard being evaluated<sup>a</sup> |  |  | Food loss (%)<br>products meeting<br>Implication<br>for the<br>contamina<br>tion profile<br>described<sup>b</sup> | Relative Risk<sup>c</sup> of Illness per Serving from Characterized Contamination Compared to Baseline Illness<sup>d</sup> |  |  |  |  |  |
| --- | --- | --- | --- | --- | --- | --- | --- | --- | --- |
| Type | Level (CFU/g) | Serotypes |  | A1<sup>e</sup> – All serotypes equal – R | A1<sup>e</sup> – All serotypes equal – @ Risk in Excel | A1.1<sup>e</sup> – All serotypes equal using triangular distribution – @ Risk in Excel | A2<sup>e</sup> – Reduced virulence Kentucky – R | A2<sup>e</sup> – Reduced virulence Kentucky – @ Risk in Excel | A2.1<sup>e</sup> – Reduced virulence for Kentucky using Anatum Dr model and WHO/FAO (2002) using triangular distribution – @ Risk in Excel |
| Baseline | All (No threshold) | All | 100% | 100.0% | 100.0% | 100.0% | 100.0% | 100.0% | 100.0% |
| Level only | >0.0074 | All | 20.7% | 98.4% | 98.3% | 98.3% | 98.4% | 98.4% | 98.2% |
|  | >1 | All | 0.67% | 68.1% | 67.3% | 66.8% | 68.0% | 68.5% | 65.1% |
|  | >10 | All | 0.059% | 37.4% | 36.0% | 35.6% | 37.0% | 41.9% | 34.3% |
| Level & high-virulent (KPI)<sup>f</sup> serotypes | >0.0074 | Enteritidis, Infantis, or Typhimurium | 10.6% | 50.2% | 47.3% | 50.6% | 70.2% | 73.8% | 63.5% |
|  | >1 | Enteritidis, Infantis, or Typhimurium | 0.34% | 34.7% | 31.4% | 34.6% | 48.4% | 52.1% | 39.9% |
|  | >10 | Enteritidis, Infantis, or Typhimurium | 0.029% | 19.1% | 16.0% | 20.1% | 26.3% | 32.4% | 18.4% |
| Level & exclude low-virulent serotype | >0.0074 | All except Kentucky | 14.6% | 68.7% | 66.2% | 71.5% | 97.3% | 97.1% | 97.4% |
|  | >1 | All except Kentucky | 0.48% | 47.2% | 44.2% | 49.7% | 67.3% | 67.5% | 64.5% |
|  | >10 | All except Kentucky | 0.041% | 25.5% | 22.2% | 27.9% | 36.6% | 41.2% | 34.0% |

<sup>a</sup> Three candidate final product standards are shown on the left side of the table. On the right side of the standards, the relative risk from two different *Salmonella* level distribution inputs.
<sup>b</sup> Proportion of products meeting the described contamination profile was collected from a simulation from A3.
<sup>c</sup> Relative Risk of Illness per Serving (%) is calculated as: The mean number of illnesses estimated from products meeting characterized contamination/The mean number of illnesses estimated from all products x 100. The relative risk is calculated by running one simulation of 1 million iterations in R. The baseline probability of illness in a serving was scaled to 2 illnesses in a million servings.
<sup>d</sup> Different scaling factors were used for each approach and the dataset to match the baseline probability of illness per serving to about 1.9 in a million ( $1.9 \times 10^{-6}$ illness/serving) based on calculations from public health data (Refer to Simulating Salmonella dose in one serving using a scaling factor to adjust for retail to consumer handling practices in materials and methods section). The scaling factor accounts for all customer and consumer handling from packing the finished product to consumption. Baseline mean illness was calculated by multiplying this baseline probability of illness per serving and the total number of servings in a year.
<sup>e</sup> A1: All serotypes equal; A1.1: All serotypes equal using triangular distribution for DR parameters; A2: Reduced virulence for Kentucky using Anatum DR model, A2.1: Reduced virulence for Kentucky using Anatum DR model and WHO/FAO (2002) using triangular distribution for DR parameters.
<sup>f</sup> KPI serotype is a Key Performance Indicator serotype which was defined by USDA-FSIS for poultry as Enteritidis, Infantis and Typhimurium at the time of this work.

**Table S3.**
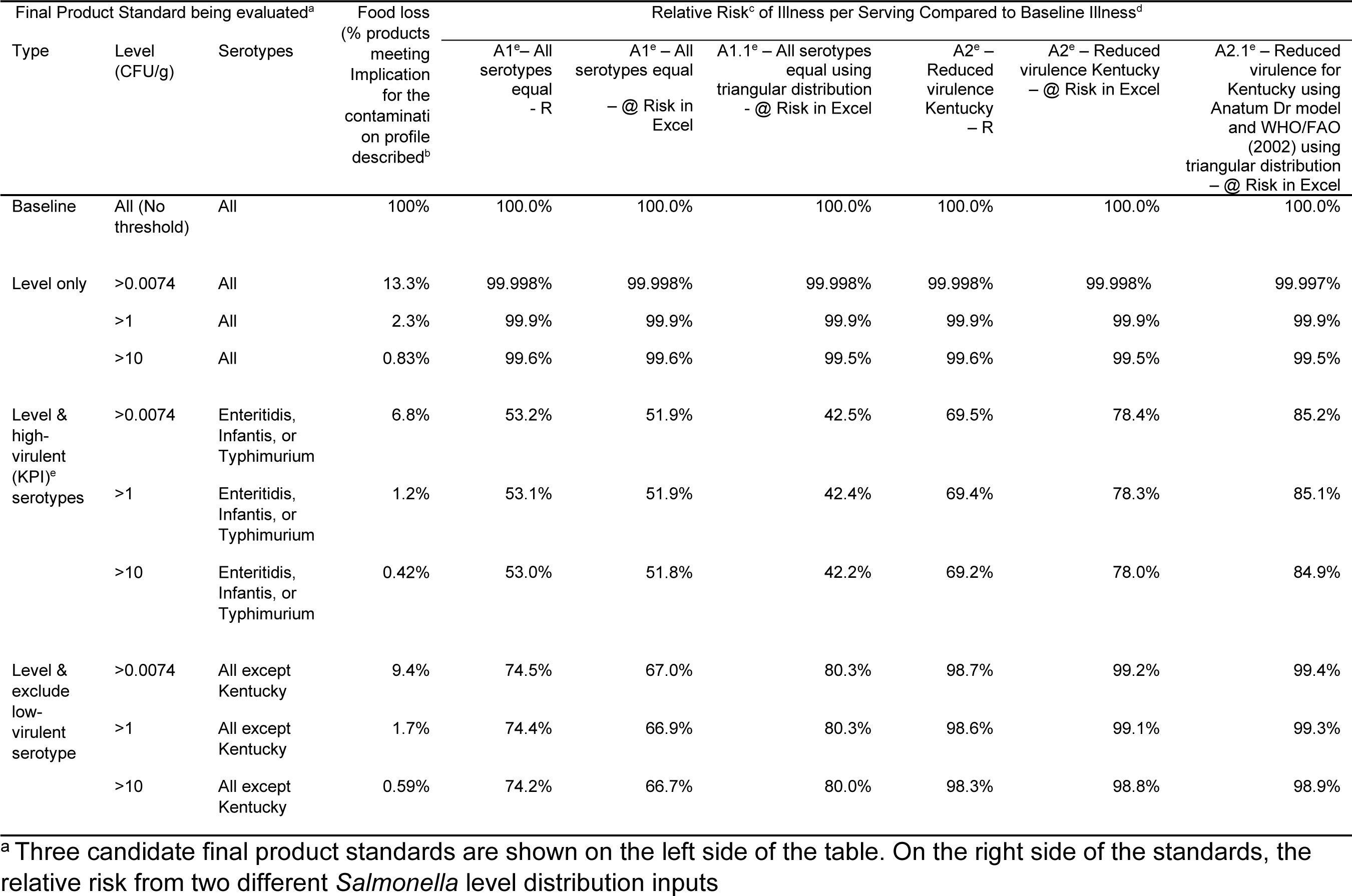
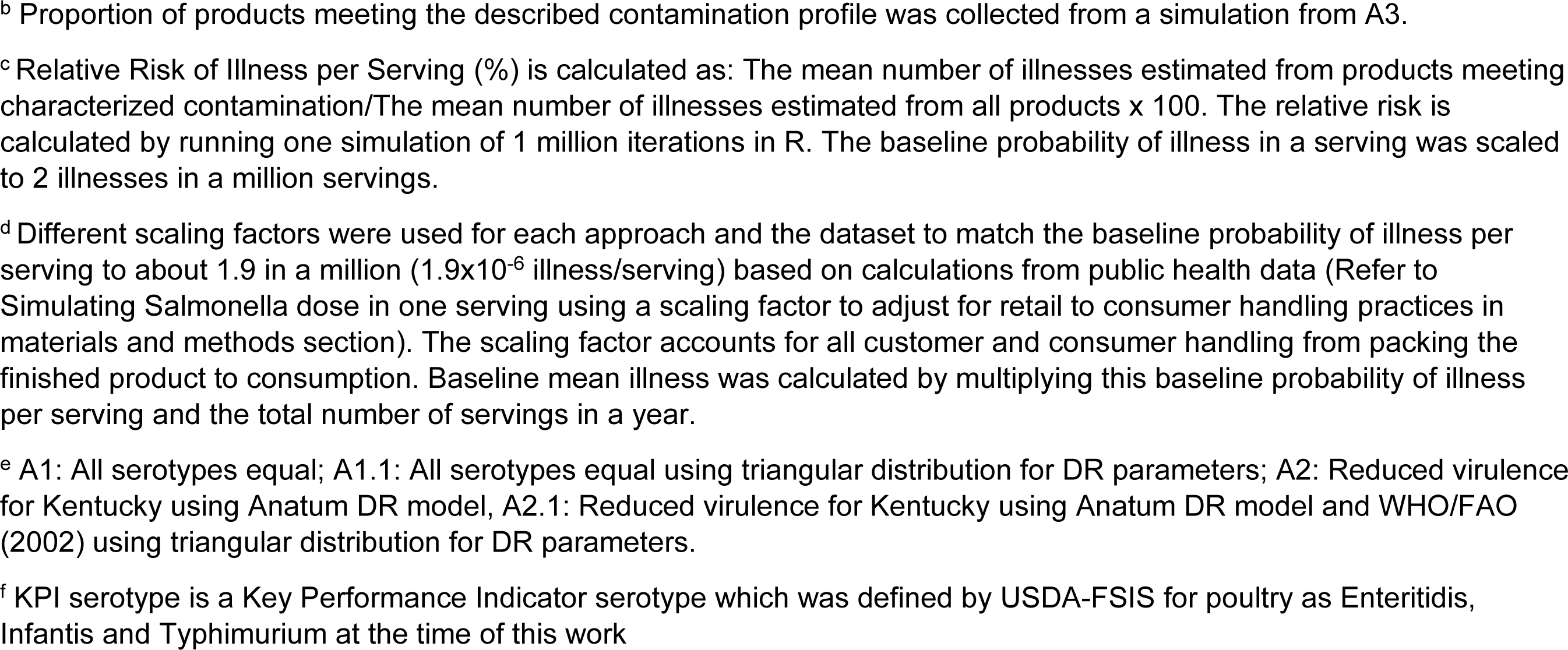
Comparing relative risk of illnesses for the implementation of models in R or @RISK for Excel from 2023 HACCP verification data input level.

| Final Product Standard being evaluated<sup>a</sup> |  |  |  | Relative Risk<sup>c</sup> of Illness per Serving Compared to Baseline Illness<sup>d</sup> |  |  |  |  |  |
| --- | --- | --- | --- | --- | --- | --- | --- | --- | --- |
| Type | Level (CFU/g) | Serotypes | Food loss (% products meeting Implication for the contamination on profile described<sup>b</sup>) | A1<sup>e</sup> – All serotypes equal – R | A1<sup>e</sup> – All serotypes equal – @ Risk in Excel | A1.1<sup>e</sup> – All serotypes equal using triangular distribution – @ Risk in Excel | A2<sup>e</sup> – Reduced virulence Kentucky – R | A2<sup>e</sup> – Reduced virulence Kentucky – @ Risk in Excel | A2.1<sup>e</sup> – Reduced virulence for Kentucky using Anatum Dr model and WHO/FAO (2002) using triangular distribution – @ Risk in Excel |
| Baseline | All (No threshold) | All | 100% | 100.0% | 100.0% | 100.0% | 100.0% | 100.0% | 100.0% |
| Level only | >0.0074 | All | 13.3% | 99.998% | 99.998% | 99.998% | 99.998% | 99.998% | 99.997% |
|  | >1 | All | 2.3% | 99.9% | 99.9% | 99.9% | 99.9% | 99.9% | 99.9% |
|  | >10 | All | 0.83% | 99.6% | 99.6% | 99.5% | 99.6% | 99.5% | 99.5% |
| Level & high-virulent (KPI)<sup>e</sup> serotypes | >0.0074 | Enteritidis, Infantis, or Typhimurium | 6.8% | 53.2% | 51.9% | 42.5% | 69.5% | 78.4% | 85.2% |
|  | >1 | Enteritidis, Infantis, or Typhimurium | 1.2% | 53.1% | 51.9% | 42.4% | 69.4% | 78.3% | 85.1% |
|  | >10 | Enteritidis, Infantis, or Typhimurium | 0.42% | 53.0% | 51.8% | 42.2% | 69.2% | 78.0% | 84.9% |
| Level & exclude low-virulent serotype | >0.0074 | All except Kentucky | 9.4% | 74.5% | 67.0% | 80.3% | 98.7% | 99.2% | 99.4% |
|  | >1 | All except Kentucky | 1.7% | 74.4% | 66.9% | 80.3% | 98.6% | 99.1% | 99.3% |
|  | >10 | All except Kentucky | 0.59% | 74.2% | 66.7% | 80.0% | 98.3% | 98.8% | 98.9% |

<sup>a</sup> Three candidate final product standards are shown on the left side of the table. On the right side of the standards, the relative risk from two different *Salmonella* level distribution inputs
<sup>b</sup> Proportion of products meeting the described contamination profile was collected from a simulation from A3.
<sup>c</sup> Relative Risk of Illness per Serving (%) is calculated as: The mean number of illnesses estimated from products meeting characterized contamination/The mean number of illnesses estimated from all products x 100. The relative risk is calculated by running one simulation of 1 million iterations in R. The baseline probability of illness in a serving was scaled to 2 illnesses in a million servings.
<sup>d</sup> Different scaling factors were used for each approach and the dataset to match the baseline probability of illness per serving to about 1.9 in a million ( $1.9 \times 10^{-6}$ illness/serving) based on calculations from public health data (Refer to Simulating Salmonella dose in one serving using a scaling factor to adjust for retail to consumer handling practices in materials and methods section). The scaling factor accounts for all customer and consumer handling from packing the finished product to consumption. Baseline mean illness was calculated by multiplying this baseline probability of illness per serving and the total number of servings in a year.
<sup>e</sup> A1: All serotypes equal; A1.1: All serotypes equal using triangular distribution for DR parameters; A2: Reduced virulence for Kentucky using Anatum DR model, A2.1: Reduced virulence for Kentucky using Anatum DR model and WHO/FAO (2002) using triangular distribution for DR parameters.
<sup>f</sup> KPI serotype is a Key Performance Indicator serotype which was defined by USDA-FSIS for poultry as Enteritidis, Infantis and Typhimurium at the time of this work

## References

1. Barrow, P. A., Jones, M. A., Smith, A. L., & Wigley, P. (2012). The long view: *Salmonella* – the last forty years. Avian Pathology, 41, 413–420. 10.1080/03079457.2012.718071

2. Buchanan, R. L., Gorris, L. G. M., Hayman, M. M., Jackson, T. C., & Whiting, R. C. (2017). A review of *Listeria monocytogenes*: An update on outbreaks, virulence, dose-response, ecology, and risk assessments. Food Control, 75, 1–13. 10.1016/j.foodcont.2016.12.016

3. CDC. (2018). National Health and Nutrition Examination Survey: NHANES 2013-2014. Centers for Disease Control and Prevention. https://wwwn.cdc.gov/nchs/nhanes/search/datapage.aspx?Component=Dietary&CycleBeginYear=2013. Accessed Feb 28, 2024.

4. CDC (2023). *Salmonella* Outbreaks Associated with Not Ready-to-Eat Breaded, Stuffed Chicken Products—United States, 1998–2022. Morbidity and Mortality Weekly Report, 72, 484. 10.15585/mmwr.mm7218a2. Accessed Feb 28, 2024.

5. Chavez-Velado, D. R., Vargas, D. A., & Sanchez-Plata, M. X. (2024). Bio-Mapping *Salmonella* and *Campylobacter* Loads in Three Commercial Broiler Processing Facilities in the United States to Identify Strategic Intervention Points. Foods, 13. 10.3390/foods13020180

6. Cheng, R. A., Eade, C. R., & Wiedmann, M. (2019). Embracing Diversity: Differences in Virulence Mechanisms, Disease Severity, and Host Adaptations Contribute to the Success of Nontyphoidal *Salmonella* as a Foodborne Pathogen. Frontiers in Microbiology, 10, 1368. 10.3389/fmicb.2019.01368

7. Cohn, A., Gremillion, T., Hedberg, C., Kincheloe, J., Robach, M., Stasiewicz, M., & Wiedmann, M. (2023). Risk Management Options to Reduce Human Salmonellosis Cases Due to Consumption of Raw Poultry. Food Protection Trends, 43. 10.4315/FPT-22-035

8. Cosby, D. E., Cox, N. A., Harrison, M. A., Wilson, J. L., Buhr, R. J., & Fedorka-Cray, P. J. (2015). *Salmonella* and antimicrobial resistance in broilers: A review. Journal of applied poultry research, 24, 408–426. 10.3382/japr/pfv038

9. Delignette-Muller, M.-L., & Dutang, C. (2015). Package ’fitdistrplus’: Help to Fit of a Parametric Distribution to Non-Censored or Censored Data. In CRAN (Version 1.1–11) https://cran.r-project.org/web/packages/fitdistrplus/index.html

10. Dórea, F. C., Cole, D. J., Hofacre, C., Zamperini, K., Mathis, D., Doyle, M. P., Lee, M. D., & Maurer, J. J. (2010). Effect of *Salmonella* vaccination of breeder chickens on contamination of broiler chicken carcasses in integrated poultry operations.Applied and environmental microbiology,76, 7820–7825. 10.1128/aem.01320-10

11. Fenske, G. J., Pouzou, J. G., Pouillot, R., Taylor, D. D., Costard, S., & Zagmutt, F. J. (2023). The genomic and epidemiological virulence patterns of *Salmonella enterica* serovars in the United States. PloS one, 18, e0294624. 10.1371/journal.pone.0294624

12. Ferrari, R. G., Rosario, D. K. A., Cunha-Neto, A., Mano, S. B., Figueiredo, E. E. S., & Conte- Junior, C. A. (2019). Worldwide Epidemiology of *Salmonella* Serovars in Animal-Based Foods: a Meta-analysis. Applied and Environtal Microbiology, 85, e00591–00519. doi:10.1128/AEM.00591-19

13. Fierer, J., & Guiney, D. G. (2001). Diverse virulence traits underlying different clinical outcomes of *Salmonella* infection. Journal of Clinical Investigation, 107, 775–780. 10.1172/JCI12561

14. Foley, S. L., R. Nayak, I. B. Hanning, T. J. Johnson, J. Han, & S. C. Ricke. (2011). Population Dynamics of *Salmonella enterica* Serotypes in Commercial Egg and Poultry Production. Applied and Environtal Microbiology, 77:4273–4279. 10.1128%2FAEM.00598-11

15. Grimont, P., & Weill, F. (2007). WHO collaborating centre for reference and research on *Salmonella*. Antigenic formulae of the Salmonella serovars, 6-10. https://www.researchgate.net/publication/283428414_Antigenic_Formulae_of_the_Salmonella_serovars_9th_ed_Paris_WHO_Collaborating_Centre_for_Reference_and_Research_on_Salmonella

16. Grosjean, P., & Ibanez, F. (2018). Package ’pastecs’: Package for Analysis of Space-Time Ecological Series. In CRAN (Version 1.3.21) https://CRAN.R-project.org/package=pastecs

17. Hofacre, C. L., Rosales, A. G., Da Costa, M., Cookson, K., Schaeffer, J., & Jones, M. K. (2021). Immunity and protection provided by live modified vaccines against paratyphoid *Salmonella* in poultry—an applied perspective. Avian Diseases, 65, 295–302. 10.1637/aviandiseases-D-20-00126

18. Jackson, B. R., Griffin, P. M., Cole, D., Walsh, K. A., & Chai, S. J. (2013). Outbreak-associated *Salmonella enterica* serotypes and food Commodities, United States, 1998-2008. Emerging Infectous Diseases, 19, 1239-1244. 10.3201/eid1908.121511

19. Jones, T. F., Ingram, L. A., Cieslak, P. R., Vugia, D. J., Tobin-D’Angelo, M., Hurd, S., Medus, C., Cronquist, A., & Angulo, F. J. (2008). Salmonellosis outcomes differ substantially by serotype. Journal of Infectious Diseases, 198, 109–114. 10.1086/588823

20. Jongenburger, I., Den Besten, H. M. W., & Zwietering, M. H. (2015). Statistical aspects of food safety sampling. Annual Review Food Science and Technology, 6, 479–503. 10.1146/annurev-food-022814-015546

21. Kim, S. A., Park, S. H., Lee, S. I., & Ricke, S. C. (2017). Development of a rapid method to quantify *Salmonella* Typhimurium using a combination of MPN with qPCR and a shortened time incubation. Food Microbiology, 65, 7–18. 10.1016/j.fm.2017.01.013

22. Lambertini, E., Ruzante, J. M., Chew, R., Apodaca, V. L., & Kowalcyk, B. B. (2019). The public health impact of different microbiological criteria approaches for *Salmonella* in chicken parts. Microbial Risk Analysis, 12, 44–59. 10.1016/j.mran.2019.06.002

23. Lambertini, E., Ruzante, J. M., & Kowalcyk, B. B. (2021). The Public Health Impact of Implementing a Concentration-Based Microbiological Criterion for Controlling *Salmonella* in Ground Turkey. Risk Analysis, 41, 1376–1395. 10.1111/risa.13635

24. Luvsansharav, U. O., Vieira, A., Bennett, S., Huang, J., Healy, J. M., Hoekstra, R. M., Bruce, B. B., & Cole, D. (2019). *Salmonella* Serotypes: A Novel Measure of Association with Foodborne Transmission. Foodborne Pathogens and Diseases, 17, 151–155. 10.1089/fpd.2019.2641

25. Mataragas, M., Drosinos, E. H., Tsola, E., & Zoiopoulos, P. E. (2012). Integrating statistical process control to monitor and improve carcasses quality in a poultry slaughterhouse implementing a HACCP system. Food Control, 28, 205–211. 10.1016/j.foodcont.2012.05.032

26. Miller, R. A., & Wiedmann, M. (2016). The cytolethal distending toxin produced by nontyphoidal *Salmonella* serotypes Javiana, Montevideo, Oranienburg, and Mississippi induces DNA damage in a manner similar to that of serotype Typhi. MBio, 7, 10–1128. 10.1128/mbio.02109-16

27. NACMCF. (2024). Response to Questions Posed by the Food Safety and Inspection Service: Enhancing *Salmonella* Control in Poultry Products. Journal of Food Protection, 87, 100168. 10.1016/j.jfp.2023.100168

28. Nauta, M. J., van der Wal, F. J., Putirulan, F. F., Post, J., van de Kassteele, J., & Bolder, N. M. (2009). Evaluation of the “testing and scheduling” strategy for control of *Campylobacter* in broiler meat in The Netherlands. International Journal of Food Microbiology, 134, 216–222. 10.1016/j.ijfoodmicro.2009.06.014

29. Plummer, M. (2022). JAGS: Just Another Gibbs Sampler. In (Version 4.3.1) https://mcmc-jags.sourceforge.io/

30. Plummer, M., Stukalov, A., & Denwood, M. (2023). Package‘rjags’: Interface to the JAGS MCMC library. In CRAN https://cran.r-project.org/web/packages/rjags/rjags.pdf

31. Office of Disease Prevention and Health Promotion. (2023). US Department of Health and Human Services: Healthy People 2030. https://health.gov/healthypeople/objectives-and-data/browse-objectives/foodborne-illness. Accessed Feb 28, 2024.

32. R Core Team. (2023). R: A Language and Environment for Statistical Computing. In (Version 4.3.1) R Foundation for Statistical Computing. https://www.r-project.org/

33. Rasschaert, G., Houf, K., Godard, C., Wildemauwe, C., Pastuszczak-Frak, M., & De Zutter, L. (2008). Contamination of Carcasses with *Salmonella* during Poultry Slaughter. Journal of Food Protection, 71, 146–152. 10.4315/0362-028X-71.1.146

34. Rasschaert, G., L. De Zutter, L. Herman, and M. Heyndrickx. (2020). *Campylobacter* contamination of broilers: the role of transport and slaughterhouse. International Journal of Food Microbiology, 10.1016/j.ijfoodmicro.2020.108564

35. Richards, A. K., Kue, S., Norris, C. G., & Shariat, N. W. (2023). Genomic and phenotypic characterization of *Salmonella enterica* serovar Kentucky. Microbial Genomics, 9. 10.1099/mgen.0.001089

36. Richards, A. K., Siceloff, A. T., Simmons, M., Tillman, G. E., & Shariat, N. W. (2023). Poultry processing interventions reduce *Salmonella* serovar complexity on post-chill young chicken carcasses as determined by deep serotyping. Journal of Food Protection, 100208. 10.1016/j.jfp.2023.100208

37. Ripley, B., & Venables, B. (2023). Package ’MASS’: Support Functions and Datasets for Venables and Ripley’s MASS. In CRAN (Version 7.3–60) https://cran.r-project.org/web/packages/MASS/index.html

38. Sampedro, F., Garcés-Vega, F., Strickland, A. J., & Hedberg, C. W. (2024). Developing a risk management framework to improve public health outcomes by enumerating and serotyping *Salmonella* in ground turkey. Epidemiology & Infectection, 152, e12. 10.1017/s0950268823002029

39. Scallan, E., Hoekstra, R. M., Angulo, F. J., Tauxe, R. V., Widdowson, M. A., Roy, S. L., Jones, J. L., & Griffin, P. M. (2011). Foodborne illness acquired in the United States--major pathogens. Emerging Infectious Diseases, 17, 7–15. 10.3201/eid1701.p11101

40. Siceloff, A. T., Waltman, D., & Shariat, N. W. (2022). Regional *Salmonella* Differences in United States Broiler Production from 2016 to 2020 and the Contribution of Multiserovar Populations to *Salmonella* Surveillance. Applied and Environtal Microbiology, 88, e0020422. 10.1128/aem.00204-22

41. Strickland, A. J., Sampedro, F., & Hedberg, C. W. (2023). Quantitative Risk Assessment of *Salmonella* in Ground Beef Products and the Resulting Impact of Risk Mitigation Strategies on Public Health. Journal of Food Protection, 86, 100093. 10.1016/j.jfp.2023.100093

42. Tate, H., C. H. Hsu, J. C. Chen, J. Han, S. L. Foley, J. P. Folster, L. K. Francois Watkins, J. Reynolds, G. E. Tillman, E. Nyirabahizi, and S. Zhao. (2022). Genomic Diversity, Antimicrobial Resistance, and Virulence Gene Profiles of *Salmonella* Serovar Kentucky Isolated from Humans, Food, and Animal Ceca Content Sources in the United States. Foodborne Pathogens and Diseases. 19, 509–521. 10.1089/fpd.2022.0005

43. Teunis, P. F., & Havelaar, A. H. (2000). The Beta Poisson dose-response model is not a single- hit model. Risk Analysis, 20, 513–520. 10.1111/0272-4332.204048

44. Teunis, P. F. M. (2022). Dose response for *Salmonella* Typhimurium and Enteritidis and other nontyphoid enteric *salmonellae*. Epidemics, 41, 100653. 10.1016/j.epidem.2022.100653

45. Teunis, P. F. M., Kasuga, F., Fazil, A., Ogden, I. D., Rotariu, O., & Strachan, N. J. C. (2010). Dose–response modeling of *Salmonella* using outbreak data. International Journal of Food Microbiology, 144, 243–249. 10.1016/j.ijfoodmicro.2010.09.026

46. The Interagency Food Safety Analytics Collaboration. (2022). Foodborne illness source attribution estimates for 2019 for *Salmonella*, *Escherichia coli* O157, *Listeria monocytogenes*, and *Campylobacter* using multi-year outbreak surveillance data, United States. https://www.cdc.gov/foodsafety/ifsac/pdf/P19-2019-report-TriAgency-508.pdf. Accessed Feb 28, 2024.

47. The Interagency Food Safety Analytics Collaboration. (2023). Foodborne illness source attribution estimates for 2019 for *Salmonella*, *Escherichia coli* O157, *Listeria monocytogenes*, and *Campylobacter* using multi-year outbreak surveillance data, United States. https://www.cdc.gov/foodsafety/ifsac/pdf/P19-2019-report-TriAgency-508.pdf. Accessed Feb 28, 2024.

48. Thompson, C. P., Doak, A. N., Amirani, N., Schroeder, E. A., Wright, J., Kariyawasam, S., Lamendella, R., & Shariat, N. W. (2018). High-Resolution Identification of Multiple *Salmonella* Serovars in a Single Sample by Using CRISPR-SeroSeq. Applied and Environmental Microbiology, 84, e01859–01818. doi:10.1128/AEM.01859-18

49. United States Census Bureau. (2023). U.S. and World Population Clock. Retrieved 2023-11-7 from https://www.census.gov/popclock/. Accessed Feb 28, 2024.

50. USDA-ERS. (2023). Food Availability and Consumption. https://www.ers.usda.gov/data-products/ag-and-food-statistics-charting-the-essentials/food-availability-and-consumption/. Accessed Feb 28, 2024.

51. USDA-FSIS. (2012). The nationwide microbiological baseline data collection program: raw chicken parts survey, January 2012–August 2012. Food Safety and Inspection Service, US Department of Agriculture, Washington, DC. , 28. https://www.fsis.usda.gov/sites/default/files/media_file/2020-07/Baseline_Data_Raw_Chicken_Parts.pdf. Accessed Feb 28, 2024.

52. USDA-FSIS. (2014). MLG Appendix 2.05: Most Probable Number Procedure and Tables. https://www.fsis.usda.gov/sites/default/files/media_file/2021-03/MLG-Appendix-2.pdf. Accessed Feb 28, 2024.

53. USDA-FSIS. (2016). New performance standards for *Salmonella* and *Campylobacter* in not- ready-to-eat comminuted chicken and turkey products and raw chicken parts and changes to related agency verification procedures: Response to comments and announcement of implementation schedule. Fed Reg, 81, 7285–7300.

54. USDA-FSIS. (2022). Proposed Framework for Controlling *Salmonella* in Poultry. Fed Reg, 87, 62784–62786.

55. USDA-FSIS. (2023a). FY 2022-2026 Food Safety Key Performance Indicator. Retrieved from https://www.fsis.usda.gov/inspection/inspection-programs/inspection-poultry-products/reducing-salmonella-poultry/salmonella-0. Accessed Feb 28, 2024.

56. USDA-FSIS. (2023b). Laboratory Sampling Data. Retrieved from https://www.fsis.usda.gov/science-data/data-sets-visualizations/laboratory-sampling-data. Accessed Feb 28, 2024.

57. USDA-FSIS. (2023c). MLG 4.14 Isolation and Identification of Salmonella from Meat, Poultry, Pasteurized Egg, Siluriformes (Fish) Products and Carcass and Environmental Sponges Retrieved from https://www.fsis.usda.gov/sites/default/files/media_file/documents/MLG-4.14.pdf. Accessed Feb 28, 2024.

58. USDA-FSIS. (2023d). *Salmonella* By the Numbers. https://www.fsis.usda.gov/inspection/inspection-programs/inspection-poultry-products/reducing-salmonella-poultry/salmonella. Accessed Feb 28, 2024.

59. USDA-FSIS. (2023e). *Salmonella* in Not-Ready-To-Eat Breaded Stuffed Chicken Products. Fed Reg, 88, 26249–26271.

60. WHO/FAO. (2002). Risk assessments of *Salmonella* in eggs and broiler chickens. Food & Agriculture Org.

61. Wickham, H. (2022). Package ’stringr’: Simple, Consistent Wrappers for Common String Operations. In CRAN (Version 1.5.0) https://CRAN.R-project.org/package=stringr

62. Wickham, H., & Bryan, J. (2023). Package ’readxl’: Read Excel Files. In CRAN (Version 1.4.3) https://cran.r-project.org/web/packages/readxl/index.html

63. Williams, M. S., & Ebel, E. D. (2012). Methods for fitting a parametric probability distribution to most probable number data. International Journal of Food Microbiology, 157, 251–258. 10.1016/j.ijfoodmicro.2012.05.014

64. Williams, M. S., Ebel, E. D., & Allender, H. D. (2015). Industry-level changes in microbial contamination on market hog and broiler chicken carcasses between two locations in the slaughter process. Food Control, 51, 361–370. 10.1016/j.foodcont.2014.11.039

65. Williams, M. S., Ebel, E. D., & Golden, N. J. (2017). Using indicator organisms in performance standards for reducing pathogen occurrence on beef carcasses in the United States. Microbial Risk Analysis, 6, 44–56. 10.1016/j.mran.2017.01.001

66. Williams, M. S., Ebel, E. D., Golden, N. J., Saini, G., Nyirabahizi, E., & Clinch, N. (2022). Assessing the effectiveness of performance standards for *Salmonella* contamination of chicken parts. International Journal of Food Microbiology, 378, 109801. 10.1016/j.ijfoodmicro.2022.109801

67. Xie, Y. (2023). Package ’knitr’: A General-Purpose Package for Dynamic Report Generation in R. In CRAN https://cran.r-project.org/web/packages/knitr/index.html

